# Proteomics-based determination of double stranded RNA interactome reveals known and new factors involved in Sindbis virus infection

**DOI:** 10.1101/2021.03.09.434591

**Authors:** Erika Girardi, Mélanie Messmer, Paula Lopez, Aurélie Fender, Johana Chicher, Béatrice Chane-Woon-Ming, Philippe Hammann, Sébastien Pfeffer

## Abstract

Viruses are obligate intracellular parasites, which depend on the host cellular machineries to replicate their genome and complete their infectious cycle. Long double stranded (ds)RNA is a common viral by-product originating during RNA virus replication and is universally sensed as a danger signal to trigger the antiviral response. As a result, viruses hide dsRNA intermediates into viral replication factories and have evolved strategies to hijack cellular proteins for their benefit. The characterization of the host factors associated with viral dsRNA and involved in viral replication remains a major challenge to develop new antiviral drugs against RNA viruses. Here, we performed anti-dsRNA immunoprecipitation followed by mass spectrometry analysis to fully characterize the dsRNA interactome in Sindbis virus (SINV) infected human cells. Among the identified proteins, we characterized SFPQ (Splicing factor, proline-glutamine rich) as a new dsRNA-associated proviral factor upon SINV infection. We showed that SFPQ depletion reduces SINV infection in human HCT116 and SK-N-BE(2) cells, suggesting that SFPQ enhances viral production. We demonstrated that the cytoplasmic fraction of SFPQ partially colocalizes with dsRNA upon SINV infection. In agreement, we proved by RNA-IP that SFPQ can bind dsRNA and viral RNA. Furthermore, we showed that overexpression of a wild type, but not an RNA binding mutant SFPQ, increased viral infection, suggesting that RNA binding is essential for its positive effect on the virus. Overall, this study provides the community with a compendium of dsRNA-associated factors during viral infection and identifies SFPQ as a new proviral dsRNA binding protein.

## Introduction

Because of their strict dependency on the host cell machinery to replicate, viruses interact with many host factors to ensure the efficient and successful completion of their life cycle. The least complex the viral genome is, the more the virus will rely on cellular proteins. For instance, positive single stranded (ss)RNA viral genomes, which mimic cellular mRNA features, typically encode for a handful of proteins allowing them to replicate and package their genome. As a consequence, these viruses are known to hijack host factors, such as regulatory non-coding RNAs (Damas et al. 2019) as well as several cellular RNA Binding Proteins (RBPs) (Nagy and Pogany 2012; Girardi et al. 2020) causing profound alterations in the cell.

Canonical RBPs participate in every step of virus infection, from the recognition of the viral RNA to its replication and translation (Nagy and Pogany 2012). In particular, a broad range of RNA viruses are able to hijack nuclear shuttling host RBPs and to alter their roles to support their own replication, thereby indirectly impacting host gene expression (Lloyd 2015). On the other hand, the cell evolved an arsenal of antiviral proteins able to recognize foreign viral features, such as non-self nucleic acids, to trigger a potent innate immune response and counteract the infection. The presence of viral double stranded (ds)RNA structures longer than 30 base pair (bp) constitutes a universal danger signal in higher eukaryotes. In mammals, the recognition of long viral dsRNA, which accumulates invariably and specifically as a by-product of viral replication or transcription, culminates with the induction of type I Interferons (IFNs) and of hundreds of interferon-stimulated-genes (ISGs). Such a chain of events perturbs the viral replication but also promotes the inhibition of protein synthesis, cellular RNA degradation and, ultimately, cell death (Stetson and Medzhitov 2006). Beside the well characterized dsRNA sensors belonging to the RIG-I like receptor (RLR) family as well as antiviral dsRNA Binding Proteins (dsRBPs) such as the protein kinase R PKR (Schlee and Hartmann 2016), additional cellular RBPs are also emerging as general or virus-specific players in the recognition and immune defence to viral dsRNA (Hur 2019). Therefore, the identification of such host factors is crucial to better understand dsRNA sensing in viral infections and to study the interactions between the host and its viruses. In particular, the importance of nuclear RBPs in the regulation of innate immune response against cytoplasmic viral RNA molecules remains underexplored.

System-wide approaches to discover the cellular RBP interactome of viral RNA are emerging as keys to deepen our understanding of the involvement of host proteins in infected cells (Garcia-Moreno et al. 2018). In particular, many efforts have been made in order to identify host factors able to bind to the viral genome (LaPointe et al. 2018; Varjak et al. 2013) and to elucidate the role of these factors at different stages of the viral life cycle. Recently, RNA interactome capture (RNA-IC) has also been employed to identify the cellular RNA-binding proteome associated with viral RNA and found out several RBPs which may regulate viral replication and could be targeted to modulate the infection (Garcia-Moreno et al. 2019; Kamel et al. 2021). However, these approaches are mainly focused on genomic or messenger viral RNAs rather than viral dsRNA replication intermediates, which are one of the key features triggering the cell immune response.

To address this lack of knowledge, we turned to an approach specifically designed to determine the dsRNA-associated proteome during viral infection. As a model for this work, we used Sindbis virus (SINV), which is a member of the alphavirus genus from the *Togaviridae* family. SINV has a capped and polyadenylated single stranded, positive-sense RNA genome of 12 kb, which consists of two open reading frames (ORFs). ORF1 encodes the non-structural polyprotein and ORF2 encodes the structural polyprotein. While the structural proteins are translated from a sub-genomic RNA, the non-structural proteins (nsP1, nsP2, nsP3 and nsP4) are directly translated from the genomic RNA since they are necessary for replication of the viral RNA inside the cytoplasm of host cells. The latter occurs through the synthesis of a complementary antigenomic RNA and the accumulation of long dsRNA intermediates in the cytoplasm of infected cells (Griffin 2007).

Here, we describe the determination of the long dsRNA interactome upon SINV infection in human HCT116 cells in an unbiased manner and aimed at characterizing the relevance of dsRNA-associated proteins during infection. To do so, we employed a novel approach based on dsRNA immunoprecipitation using a dsRNA-specific antibody followed by proteomic analysis. The outcome of the experiment revealed a number of previously known interactors but also novel factors, among which SFPQ (Splicing factor, proline-glutamine rich) that we validated as a new proviral, dsRNA-associated factor upon SINV infection.

## Results

### Identification of the dsRNA interactome in mock- and SINV-infected human cells

SINV genomic RNA replicates through the production of a reverse complementary antigenomic RNA (Figure 1A), which results in the accumulation of long dsRNA intermediates in the cytoplasm of infected cells. In order to characterize the long dsRNA interactome in mock and SINV infected human cells, we adapted the method of immunoprecipitation of long RNA duplexes (Dhir et al. 2018) using the anti-dsRNA J2 antibody, which specifically recognizes dsRNAs longer than 40 bp (Schonborn et al. 1991; Bonin et al. 2000).

**Figure 1.**
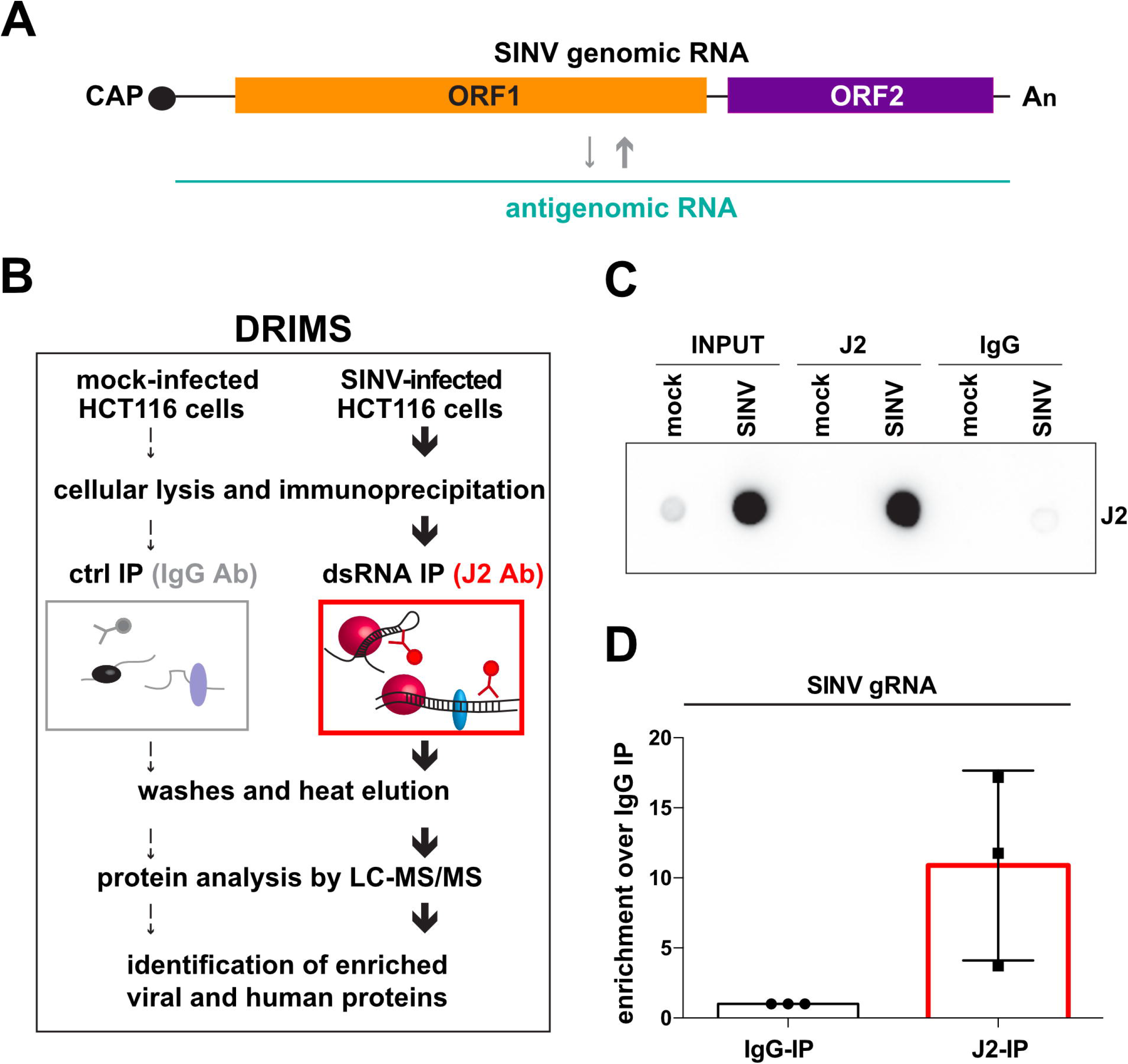
Identification of the dsRNA-associated proteome upon SINV infection in HCT116 cells. **A)** Schematic representation of SINV RNA replication and formation of dsRNA intermediates. Capped and polyadenylated viral genomic RNA is represented in black, while viral antigenomic RNA is in green. Grey arrows represent dsRNA replication. ORF1 codes for the non-structural proteins necessary for RNA replication (in orange). ORF2 codes for the structural proteins (in purple). **B**) Schematic representation of the DRIMS experimental approach. **C)** Dot blot revealed using the anti-dsRNA J2 antibody on RNA extracted from total lysate (INPUT), dsRNA-IP (J2)- or control-IP (IgG) in mock and SINV-infected HCT116 cells. **D)** RT-qPCR on SINV genomic (g)RNA upon dsRNA-IP (J2-IP) compared to control-IP (IgG-IP) on infected samples. Enrichment on three independent IP from three biological replicates are shown. Error bars represent the mean +/-standard deviation (SD).

The human colon carcinoma HCT116 cell line was chosen because it can be easily infected with SINV and is highly suitable for transfection and CRISPR-Cas9 gene editing procedures (Petitjean et al. 2020). The experimental design was based on the pull-down of long dsRNA molecules from cellular lysates of HCT116 cells infected with SINV-GFP for 24 hours at a MOI of 0.01, or mock-infected. Since we combined the anti-<u>D</u>s<u>R</u>NA Immunoprecipitation to <u>M</u>ass <u>S</u>pectrometry analysis of the isolated dsRNA-associated protein complexes, we named this approach DRIMS (Figure 1B). In order to validate the enrichment of long dsRNA molecules in SINV-infected IP conditions, we performed a dot blot analysis using the J2-antibody. We observed a specific signal both in total RNA (INPUT) and J2-IP upon SINV infection compared to the IgG-IP, suggesting that the dsRNA species accumulate in infected cells and were efficiently pulled-down by DRIMS (Figure 1C). The presence of viral RNA specifically immunoprecipitated in the infected samples was also assessed by RT-qPCR on RNA extracted from the J2- and control IgG -IP (Figure 1D).

Protein profiles in total extracts, J2- and IgG-IP eluates were analysed by silver staining of polyacrylamide gels (Figure S1A) and proteins immunoprecipitated in the J2- and IgG-IP samples were then analysed by liquid chromatography coupled to tandem mass spectrometry (LC-MS/MS). After identification and validation (FDR <= 1%) of viral and human proteins in each sample, hierarchical clustering analysis of the J2-IP replicates identified two distinct groups, indicating higher similarities in the protein profiles among the mock and infected samples, respectively (Figure S1B). Differential expression analysis performed by comparing J2-IP samples in infected *versus* mock-infected cells revealed the enrichment of both host and viral proteins upon viral infection (Figure 2A and Table S1). Of note, protein enrichment of the J2-IP over the IgG-IP samples upon SINV-GFP infection (Figure S1C and Table S2) showed similar enrichment profiles suggesting that protein binding to dsRNAs was specific. As a control, we also analysed the protein enrichment of the J2-IP over the IgG-IP samples in mock conditions, which already revealed few significantly enriched proteins, potentially associated to endogenous dsRNAs (Figure S1D and Table S3).

**Figure 2.**
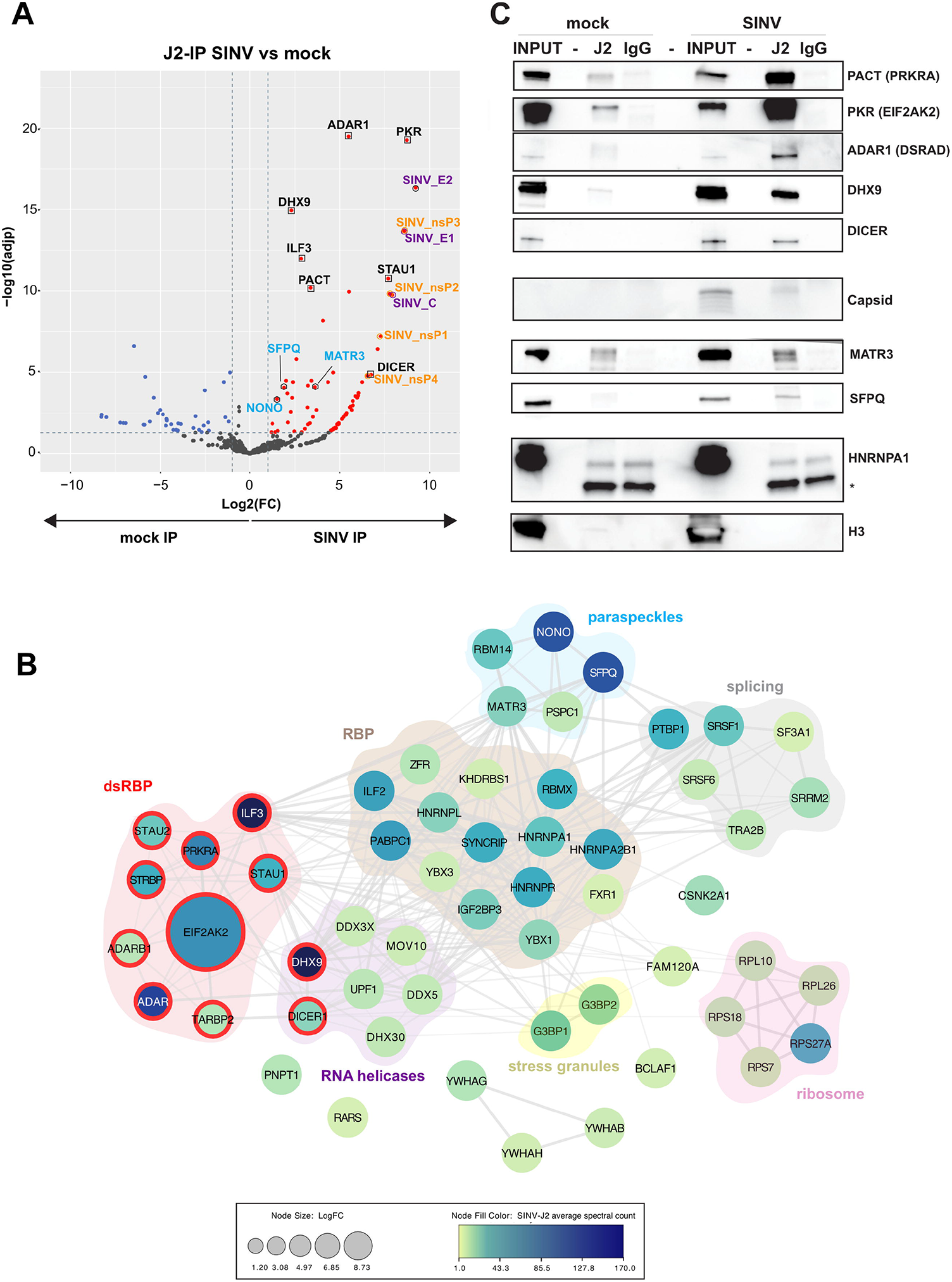
The network of dsRNA enriched proteins upon SINV infection in HCT116 cells. **A)** Volcano plot showing the global enrichment of proteins upon dsRNA-IP (J2-IP) in SINV-infected *versus* mock conditions using DRIMS data from three replicates. Infection conditions: 24 hours at MOI of 0.01. Red and blue dots represent proteins which are significantly enriched or depleted (adjusted p-value < 0.05, a minimum of 5 SpC in the most abundant condition, and abs(Log2FC) > 1), respectively, in the infected samples compared to the uninfected ones. Viral non-structural and structural proteins are indicated in orange and purple, respectively. Known dsRNA binding proteins are indicated in black and by a black square. Proteins present in paraspeckles are indicated in light blue and by a black hexagon. **B)** Visualization of the interaction network of proteins identified by DRIMS upon infection, generated with the Cytoscape StringApp. The confidence score of each interaction is mapped to the edge thickness and opacity. The size of the node relates to the enrichment in log2 fold change (LogFC) over the J2-IP mock control. The protein abundance in the SINV-J2 IP sample is illustrated by a colour scale and corresponds to the specific spectral count. Proteins were clustered into different functional categories using STRING enriched terms as a guideline. Proteins with dsRNA binding domain according to STRING enrichment (category: SMART domains) are highlighted in red. **C)** Western blots performed on total lysate (INPUT), J2- or IgG-IP in mock and SINV-GFP-infected cells using the indicated specific antibodies. Histone 3 (H3) and HNRNPA1 were used as negative controls. * corresponds to an unspecific band.

We identified a total of 63 (7 viral and 56 cellular) proteins significantly enriched in SINV-infected J2-IP samples compared to mock-infected J2-IP ones, with at least 5 spectral counts in the most abundant condition, a fold change of at least 2 and an adjusted p-value lower than 0.05 (Figure 2A and Table S1). As expected, all four viral non-structural proteins, which are necessary for SINV replication were specifically enriched in the SINV J2-IP (Figure 2A). In addition, capsid, E1 and E2 structural proteins were also immunoprecipitated. Gene Ontology (GO) term enrichment analysis of the overrepresented cellular proteins in SINV J2-IP samples based on molecular function, cellular components and biological processes indicated that most of these proteins are RNA binding proteins (RBPs) (Figure S1E, left panel), are present in both cytoplasmic and nuclear ribonucleoprotein granules (Figure S1E, middle panel) and are involved in pre-mRNA processing and mRNA metabolism (Figure S1E, right panel). Moreover, STRING analysis showed that these enriched cellular proteins mostly belong to known interaction networks (Figure 2B).

Among the cellular proteins enriched with dsRNA in infected conditions, 11 have InterPro-annotated dsRNA binding domains and are known to be involved in antiviral innate immunity to long dsRNAs. In particular, we found ADAR1 (also known as DSRAD), ADARB1, PKR (also known as EIF2AK2), PACT (also known as PRKRA), TRBP (also known as TARBP2) (Hur 2019), STAU1 (Wickham et al. 1999) and its orthologue STAU2, STRBP (Coolidge 2000), and ILF3 (Watson et al. 2019) (Figures 2A-B). Interestingly, we also identified DICER as specifically associated to dsRNAs upon SINV infection. Since the J2 Ab was previously shown to be unable to recognize pre-miRNA structures (White et al. 2014), this indicates a potential role of DICER in the recognition of long dsRNAs in SINV infected human cells. We also found several RNA helicases previously shown as regulators of antiviral innate immunity: the DExH-box RNA helicases DHX9 and DHX30 (Su et al. 2022); the DEAD-Box RNA helicases DDX3X (DEAD-Box Helicase 3 X-Linked) and DDX5 (DEAD-Box Helicase 5) (Meier-Stephenson et al. 2018); the helicases UPF1 and MOV10 (Cuevas et al. 2016). We also retrieved several RBPs previously described as having a role in SINV infection such as G3BP1 and G3BP2 (Cristea et al. 2010), which localize to stress granules, or HNRNPA1, which re-localizes to the cytoplasm upon SINV infection and enhances viral RNA translation (Lin et al. 2009) (Figures 2A-B). Many other RBPs not yet identified in the context of SINV infection were enriched in our experiments, such as PNPT1 (Dhir et al. 2018), FXR1 (Fragile X-related protein-1), ZFR (Zinc finger RNA-binding protein), TRA2B (Transformer 2 Beta Homolog), SRRM2 (Serine/Arginine Repetitive Matrix 2), SRSF6 (Serine And Arginine Rich Splicing Factor 6), RBMX (RNA-binding motif protein X-chromosome, also known as HNRNPG), KHDRBS1 (KH RNA Binding Domain Containing, Signal Transduction Associated 1), IGF2BP3 (Insulin Like Growth Factor 2 mRNA Binding Protein 3) or BCLAF (Bcl-2 Associated Transcription Factor 1). In addition, several nuclear RBPs known to be involved in paraspeckle formation, such as SFPQ (Splicing factor, proline-glutamine rich), NONO (Non-POU domain-containing octamer-binding protein), PSPC1 (Paraspeckle component 1), RBM14 (RNA Binding Motif Protein 14) and MATR3 (Matrin 3) (Nakagawa et al. 2018) showed a 1.5 to 5 log2 fold change enrichment in the SINV J2-IP compared to the mock J2-IP (Figures 2A-B). Interestingly, SFPQ and NONO were already enriched with dsRNA in uninfected conditions and their association with long dsRNA increased upon infection suggesting that they could associate with both host and viral dsRNAs (Figure S1D, Tables S1-S3).

Next, we performed western blot analyses on J2-IP and IgG-IP samples in mock and SINV-infected conditions to validate some of the enriched proteins identified by mass spectrometry (Figure 2C). We confirmed an increased association of PACT, PKR, ADAR1, DHX9 and MATR3 with dsRNA upon infection. We also validated the presence of DICER in the J2-IP only in virus-infected cells, which strengthens the potential involvement of this factor in the response to viruses in our experimental setup. Interestingly, we could confirm the enrichment of SFPQ with dsRNAs in SINV-infected cells. In contrast, our western blot analysis showed neither HNRNPA1 (a single stranded RBP) or Histone 3 (nuclear protein used as negative control) binding specifically to viral dsRNA in the J2-IP upon SINV infection (Figure 2C). Overall, our data suggest that the DRIMS approach is a powerful tool to identify long dsRNA interactome upon viral infection in human cells and, besides known antiviral factors, it allowed us to isolate several novel RBPs potentially involved in dsRNA recognition, which may affect SINV infection.

### SFPQ depletion reduces SINV infection in HCT116 and SK-N-BE(2) cells

In order to further decipher the involvement of the factors enriched with dsRNAs upon SINV infection identified by DRIMS, we performed an siRNA-based loss-of-function mini-screen on 37 candidates in HCT116 cells and quantified the effect of siRNA gene-specific treatment on SINV-GFP infection compared to the siCTRL condition by measuring GFP fluorescence intensity. As functional controls, we transfected either siRNAs directly targeting the viral 3’UTR (siSINV) (López et al. 2020) as well as siRNA targeting the antiviral protein PKR (siEIF2AK2), which reduced or increased SINV-GFP expression (Figure 3A, blue and red bars, respectively). We observed that knock-down (KD) of either *DDX5, SFPQ* or *RAN* significantly reduces GFP expression in HCT116 infected cells compared to siCTRL (Figure 3A, dark grey bars). In contrast to DDX5 and SFPQ, RAN depletion by siRNA treatment was toxic for the cells and therefore this factor was excluded from the analysis. Although the RNA helicase DDX5, which is known to play a role in other viral infections (Cheng et al. 2018), was an interesting candidate, we rather chose to pursue the functional characterisation of SFPQ in terms of its impact on SINV replication and infection. In agreement with the decrease in GFP expression (Figures 3A and B), the levels of viral capsid protein (Figure 3C), viral titers (Figure 3D) and viral genomic RNA (Figure 3E) were reduced upon *SFPQ* KD compared to the control (siCTRL). We also measured cell proliferation in siCTRL and siSFPQ treated HCT116 cells to exclude that the effect on SINV was due to siSFPQ cytotoxicity (Figure S2A). The decrease in viral capsid was also observed at a different time point (12 hpi with an MOI of 0.1) and a different MOI (24 hpi with an MOI of 0.01) (Figures S2B and C), further validating the SFPQ proviral effect. We also confirmed these findings in another cell type suitable for SINV infection, namely the SK-N-BE(2) human neuroblastoma cells (López et al. 2020), which can be differentiated into neurons upon retinoic acid (RA) treatment (Figure S2D-F). We observed not only the decrease in GFP by epifluorescence microscopy (Figure S2D), but also a reduction in the levels of viral capsid (Figure S2E) and viral titers (Figure S2F) upon *SFPQ* KD compared to the control (siCTRL).

**Figure 3.**
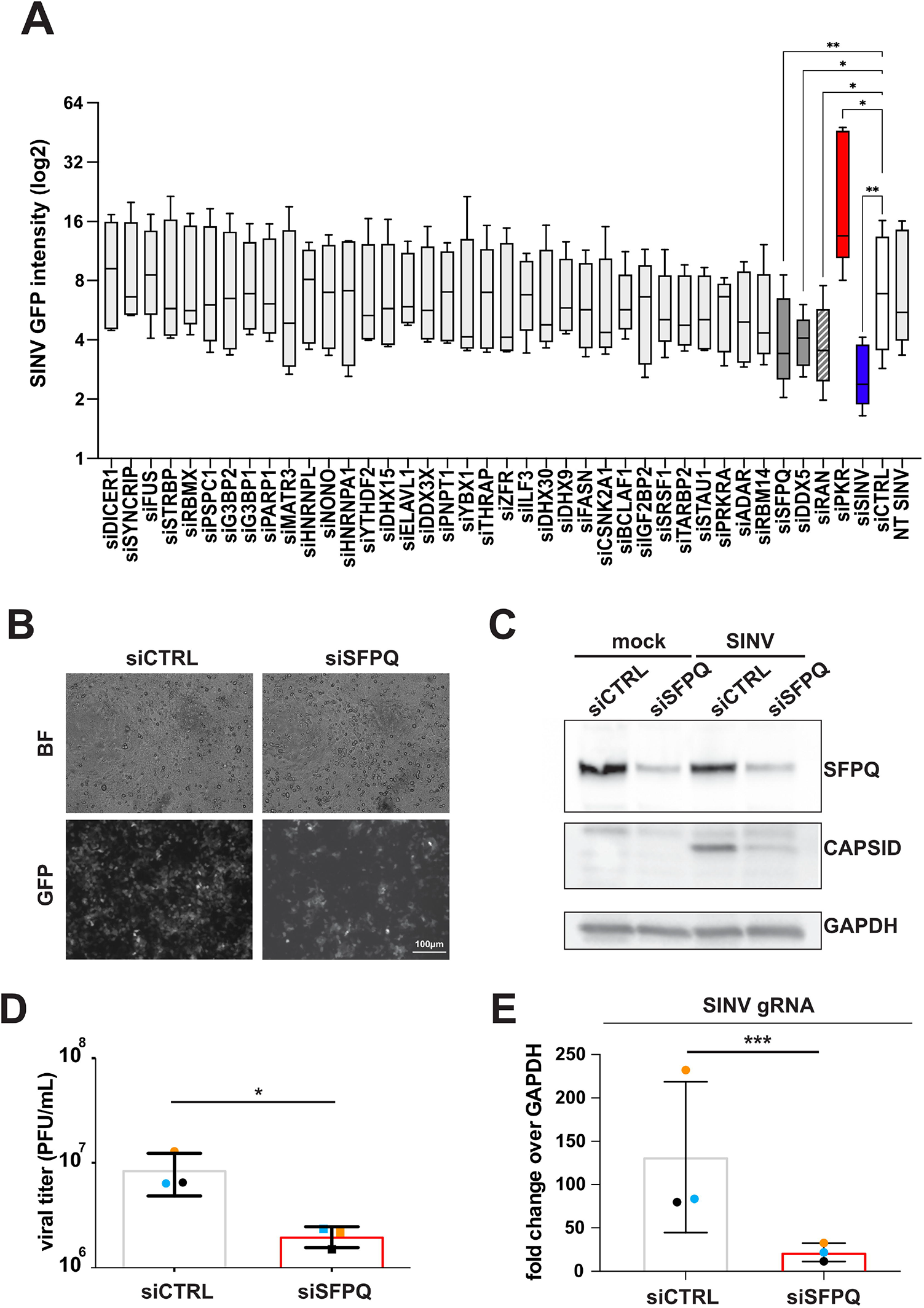
SFPQ knock-down reduces SINV-GFP infection in HCT116 cells. **A**) siRNA-based knock-down of 37 proteins enriched in SINV-J2 by DRIMS and measurement of GFP fluorescence intensity in SINV-GFP infected HCT116 cells. The GFP signal is corrected over the background fluorescence (NT mock) for each replicate and expressed in log2. SiRNAs with a significant effect on GFP accumulation relative to siCTRL (negative control, white bar) are shown in dark grey. SiRNAs against PKR mRNA (red bar) or SINV RNA (blue bar) were used as positive controls. NT SINV corresponds to not transfected (NT), SINV infected sample. Data from five independent biological replicates are shown. * p<0.05, ** p<0.01, non-parametric Friedman test with multiple comparison. **B)**. Representative pictures of SINV-GFP infected cells in siCTRL and siSFPQ treated HCT116 cells. GFP expression was measured by microscopy. BF, brightfield. Scale bar: 100 μm. **C**) Western blot performed on mock and SINV-infected cells upon siCTRL and siSFPQ treatment. Antibodies directed against SFPQ and the viral capsid protein were used. GAPDH was used as loading control. **D)** Viral production of SINV-GFP upon siCTRL and siSFPQ transfection measured by plaque assay on three independent biological replicates (PFU/mL). **E)** RT-qPCR on SINV genomic RNA relative to GAPDH upon siSFPQ treatment compared to siCTRL, in mock and SINV-GFP infected HCT116 cells. Data from three independent biological replicates are shown relative to siCTRL. Each replicate is indicated with a different coloured dot in (D) and (E). Error bars in (D) and (E) represent the mean +/-standard deviation (SD) of three independent experiments, ***p < 0.001, paired Student’s t test.

To validate the effect of *SFPQ* KD on SINV infection by an independent approach, we generated a heterozygous knock-out (KO) (+/-) *SFPQ* HCT116 cell line by CRISPR/Cas9 (Figures S3A-B). We did not manage to obtain a homozygous *SFPQ* KO cell line, probably due to the fact that SFPQ may be an essential gene (Yarosh et al. 2015). Despite the presence of one wild-type allele, we could functionally validate the negative impact of the partial SFPQ reduction on GFP expression levels (Figure S3C), capsid protein levels (Figure S3D), and viral titers (Figure S3E), in (+/-) *SFPQ* cells infected with SINV-GFP for 24 hours (MOI 0.1) compared to WT HCT116 cells. Due to its possible impact on cell viability (Yarosh et al. 2015), we measured cell proliferation in (+/-) *SFPQ* compared to WT HCT116 cells at 24, 48 and 72 hours post seeding to exclude that a defect in cell proliferation could affect our results. Indeed, we observed no effect between 24 and 48 hours, which corresponds to our experimental window of SINV infection (Figure S3F).

Overall, our results demonstrate a functional effect of SFPQ depletion on SINV infection in two independent cell types and with two different loss-of function methods, suggesting that SFPQ could play a proviral role during infection.

### SFPQ binds to cytoplasmic dsRNA and viral RNA and its RNA binding capacity is important for the proviral effect on SINV

The functional effect observed on SINV infection upon SFPQ depletion could be both dependent on and independent of SFPQ direct association to viral dsRNAs in the cytoplasm. Hence, we verified the SFPQ localization before and after SINV infection by immunofluorescence analysis (Figure 4A). As a control, the presence of long dsRNAs was observed in HCT116 cells infected with SINV for 24 hours at an MOI of 0.1 and compared to uninfected ones using the anti-dsRNA (J2) antibody (Figure S4A). Since SFPQ can interact with NONO within the paraspeckles, we also checked NONO subcellular localization and observed a nuclear accumulation in discrete foci whose intensity increased upon infection suggesting a modulation in paraspeckle composition induced by SINV in HCT116 cells (Figure S4A). We also tested the localization of G3BP1, which was immunoprecipitated together with dsRNAs by DRIMS upon infection (Figures 2A-B) and confirmed its accumulation in SINV-induced cytoplasmic stress granules (Cristea et al. 2010) (Figure S4A). In both mock and infected cells, SFPQ mostly localized to the nucleus but some signal could also be seen in the cytoplasm (Figure S4A). Such a similar SFPQ protein subcellular localization profile between mock and SINV-infected condition was also validated by nuclear/cytoplasmic fractionation experiments (Figure S4B). Nonetheless, the presence of discrete cytoplasmic SFPQ foci induced upon SINV infection suggested a change in the protein distribution within the cytoplasmic compartment specifically in the presence of the virus (Figure 4A). Co-immunofluorescence analysis showed a limited co-localization between dsRNA (antibody J2, in yellow) and SFPQ cytoplasmic granules (in pink) upon infection (Figure 4B and S4C), suggesting that a fraction of SFPQ may interact with cytoplasmic dsRNA in our experimental set-up.

**Figure 4.**
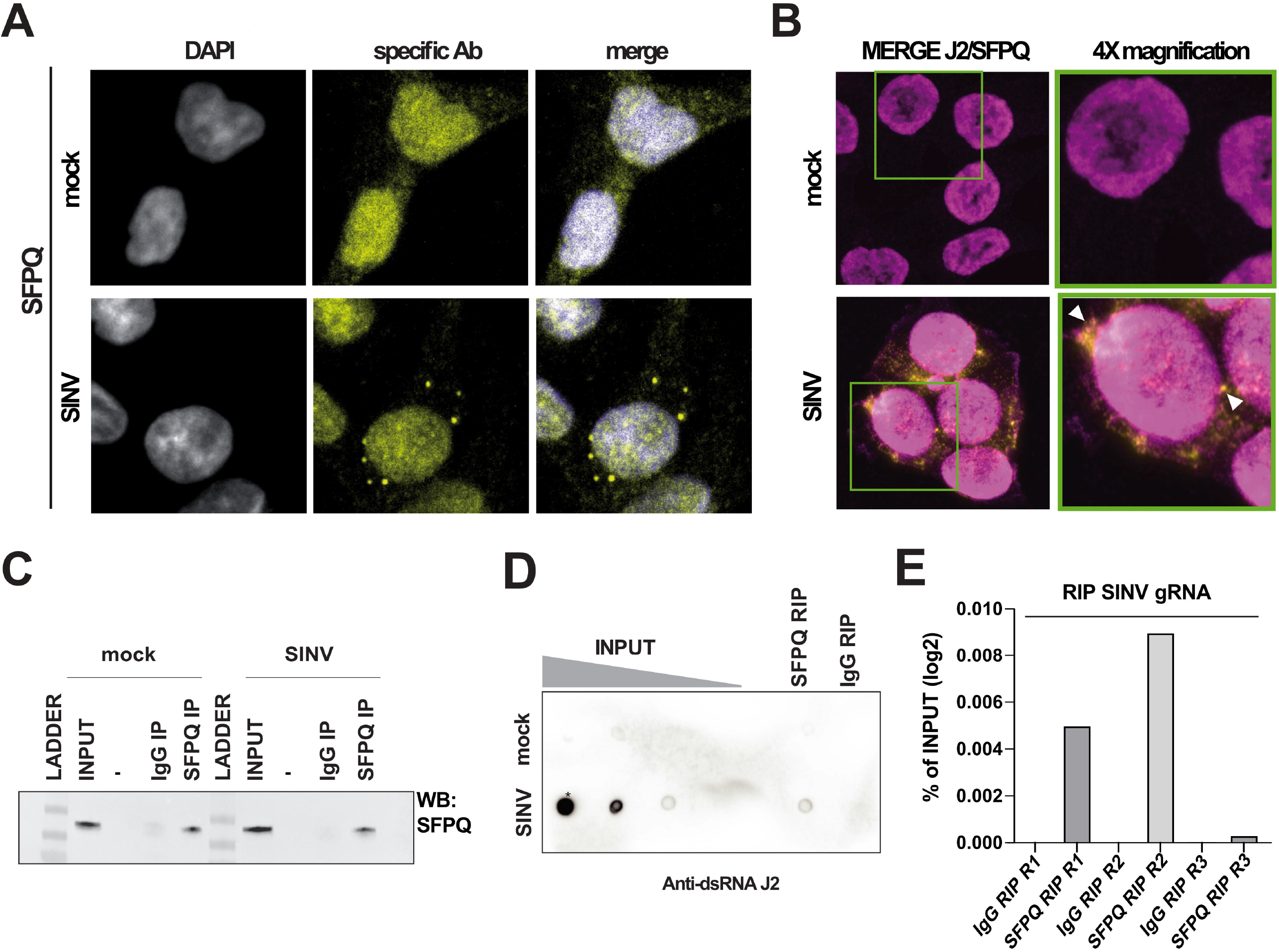
SFPQ binds to dsRNA and viral dsRNA upon SINV infection. **A**) 4X magnification of the confocal immunofluorescence images shown in Figure S4A. Anti-SFPQ mouse antibody, yellow signal. DAPI was used to stain the nuclei (in blue in the merge). **B**) Confocal co-immunofluorescence analysis on mock and SINV WT infected HCT116 cells. Merge of anti-SFPQ rabbit antibody (in purple) or anti-dsRNA J2 mouse antibody (in yellow) is shown. Single antibodies and DAPI staining are shown in Figure S4B. Magnification 63X(3X). Green squares: 4X higher magnification White arrows indicate J2 and SFPQ colocalization. **C)** SFPQ immunoprecipitation on mock and SINV-infected HCT116 samples analysed by western blot with anti-SFPQ rabbit antibody. **D)** Dot blot revealed using the anti-dsRNA J2 antibody on RNA extracted from total lysate (INPUT, four 10-fold dilutions), SFPQ- or IgG control-RIP on mock (upper part) and SINV-infected (lower part) HCT116 cells. **E)** RT-qPCR on SFPQ-IP and IgG CTRL-IP samples by using oligos to amplify SINV genomic RNA. The RNA enrichment upon SFPQ-IP or IgG IP is shown for three biological replicates and is expressed as percentage (%) of the INPUT (total RNA). The IgG RIP represent the background signal.

We further characterized the SFPQ-dsRNA interaction by immunoprecipitation of the endogenous SFPQ protein in mock and SINV infected HCT116 cells and isolation of the associated RNAs (RNA-immunoprecipitation, RIP). After verifying the SFPQ immunoprecipitation efficiency by western blot analysis (Figure 4C), we performed a dot blot analysis on immunoprecipitated RNAs to visualize long dsRNA molecules using the J2 antibody (Figure 4D). While uninfected cells did not show any dsRNA accumulation, we observed a specific signal both in the total RNA (INPUT) and the SFPQ-RIP upon SINV infection compared to the control IgG-IP, suggesting the association of dsRNA species with SFPQ in infected conditions. These data correlate with the SFPQ enrichment on dsRNAs upon SINV infection observed by DRIMS (Figures 2A-B and Table S1) and validated by western blot (Figure 2C). We also demonstrated by RT-qPCR that the viral genomic RNA was specifically enriched in the SFPQ-RIP compared to the IgG-RIP (Figure 4E) indicating that among the dsRNAs associated to SFPQ (Figure 4D) there are those of viral origin. Finally, we also proved the SFPQ association to SINV antigenomic RNA in infected conditions by semi-quantitative strand-specific RT-PCR (Figure S4D), supporting the evidence that SFPQ may indeed recognize viral dsRNA intermediates.

To rule out that SFPQ association to dsRNAs is solely mediated by its binding to flanking single stranded (ss)RNA regions, we performed J2-IP experiments followed by on-beads RNase T1 treatment (Figures S5A-B). We could confirm the efficiency of RNAse T1 digestion on total RNA (INPUT) (Figure S5A) and that the association of PKR (dsRBP, positive control) was unchanged before and after treatment, while Ago2 was not enriched (ssRBP, negative control). Interestingly, enrichment of SFPQ in the J2-IP upon infection remained unchanged before and after RNase T1 digestion, suggesting a direct binding to dsRNA regions in infected samples (Figure S5B).

Finally, to assess whether the SFPQ effect on SINV infection depends on its RNA binding capacity, we knocked-down SFPQ by transfecting a different siRNA than the previous one and directed against its 3’UTR in HCT116 cells and complemented its effect by transfecting plasmids expressing respectively myc-BFP (control), wild-type SFPQ (myc-SFPQ WT), or a mutant SFPQ deleted of both RNA Recognition Motifs (Knott et al. 2016) (myc-SFPQ ΔRRM1-2) (Figure 5A). Because the SFPQ overexpressing plasmids did not contain the 3’UTR, they were not affected by the specific siRNA treatment.

**Figure 5.**
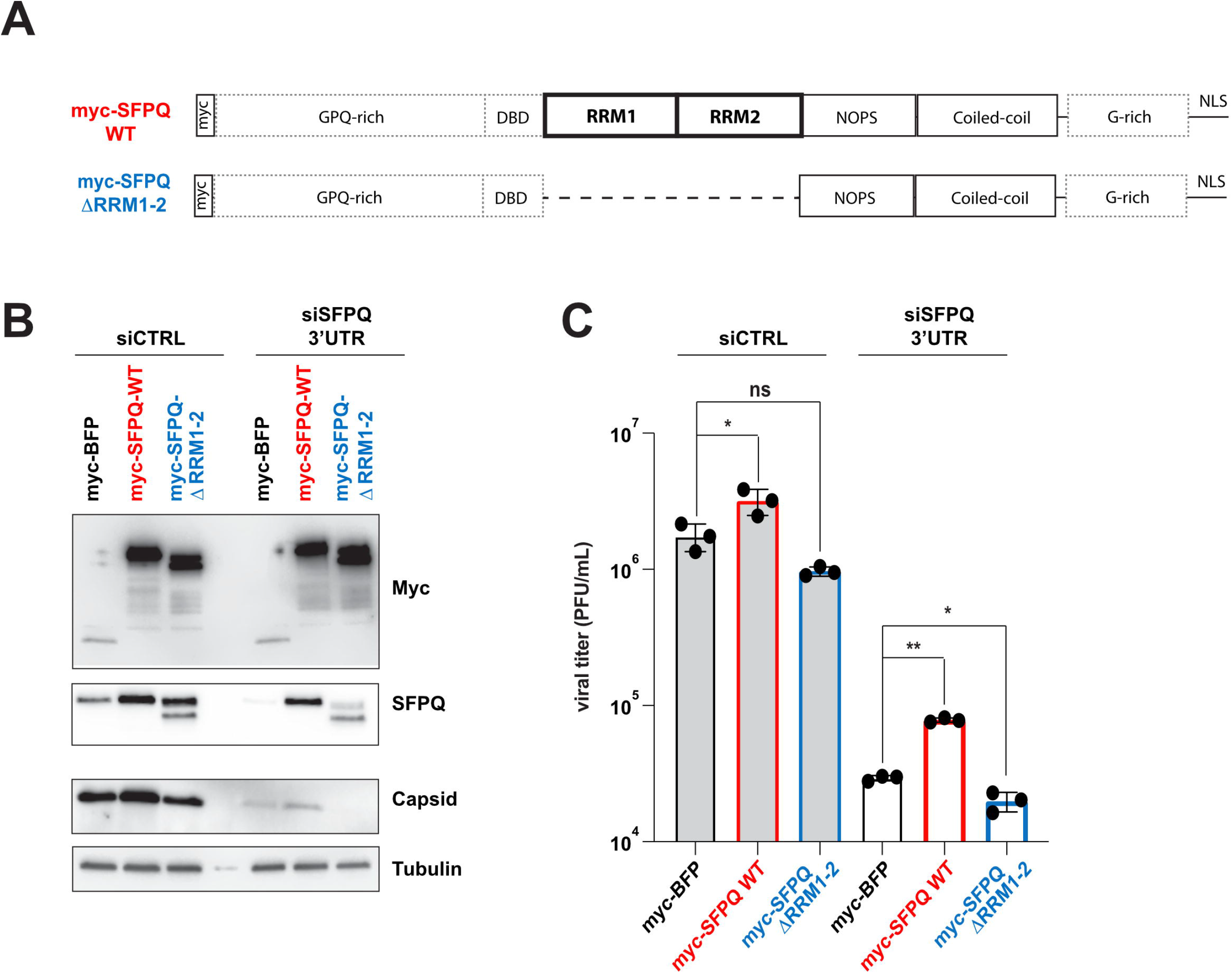
The RNA binding domains are important for the SFPQ proviral activity on SINV. **A)** Schematic representation of myc-SFPQ wild-type (WT) and ΔRRM1-2 protein architectures. The well characterized RNA recognition motifs (RRM) 1 and 2, the NonA/paraspeckle (NOPS) and the coiled-coil domains are indicated. The less characterized GPQ-rich region, the DNA-binding domain (DBD) and the G-rich region are indicated in dashed boxes (adapted from (Knott et al. 2016)). **B**) Western blot on SINV-GFP infected HCT116 cells treated with siCTRL or siSFPQ-3’UTR and then transfected with a control myc-BFP, or a myc-SFPQ WT or a myc-SFPQ ΔRRM1-2 plasmid. Antibodies directed against the myc-tag, SFPQ and the viral capsid protein were used. Tubulin was used as loading control. **C)** SINV-GFP viral production upon siRNA treatment and plasmid transfection measured by plaque assay (PFU/mL) on three independent biological replicates. Ns, non significant, * p<0.05, ** p < 0.01, One-way ANOVA with multiple comparison.

We first validated that siRNA complementary to SFPQ 3’UTR efficiently reduced endogenous SFPQ expression in HCT116 cells compared to siCTRL and that the knocked-down could be complemented by expression of the WT or ΔRRM1-2 SFPQ construct (Figure S6). Then, we measured the effect of the knock-down with or without transient expression of the SFPQ constructs. We showed that the decrease of SFPQ levels correlated with a reduction in both viral capsid expression and viral titers and that overexpression of the WT SFPQ, but not of the ΔRRM1-2 mutant, led to an increase of both viral capsid protein and viral production compared to the myc-BFP overexpression, in both siCTRL and siSFPQ-3’UTR treated cells (Figures 5B-C). However, the rescue of the knock-down by the WT SFPQ was only partial indicating that either the conditions of overexpression was suboptimal or that there are indirect effects of SFPQ knock-down that cannot be rescued efficiently by transient expression.

Overall, these findings indicate that SFPQ is able to bind viral and/or cellular RNAs or dsRNAs upon infection and that the SFPQ RNA binding domains RRM1-2 are essential for the SFPQ proviral activity on SINV.

## Discussion

Unbiased proteome-wide approaches have proven to be very useful to find new host factors regulated by or able to regulate viral infections. Over 2000 cellular RBPs have been described so far (Hentze et al. 2018), but understanding which RBP participates to the life cycle of a virus and at which step remains a challenge. To date, there has been little (Varjak et al. 2013; Incarbone et al. 2021) to no reports of the identification of proteins binding specifically to viral replication intermediates. A recent affinity proteomics approach using different synthetic nucleic acids identified novel conserved proteins able to associate to dsRNAs (Pennemann et al. 2021), but the whole picture of the viral dsRNA-associated proteome *in vivo* remains to be assessed. Here, we showed that our DRIMS approach is very effective to identify dsRNA associated proteins, since it allowed us to validate the presence of proteins with canonical dsRNA binding domains (dsRBDs) or helicase domains (Hur 2019) involved in antiviral innate immunity, such as PKR, PACT, ADAR1, STAU1, STAU2, ILF2, ILF3 and DHX9 associated to long dsRNAs upon SINV infection in human cells. Interestingly, in agreement with our recent findings (Montavon et al. 2021), we also identified DICER associated to SINV RNA duplexes. This piece of evidence further confirms the importance of DICER in modulating dsRNA-based innate immunity.

We also retrieved several RBPs which could associate with the viral duplex, even though they lack the typical dsRNA binding domains. Several examples of RBPs acting as RNA co-sensors by modulating canonical dsRNA sensor activity have been revealed so far (Liu and Gack 2020). For instance, members of the hnRNP family such as HNRNPE2 (You et al. 2009), HNRNPM (Cao et al. 2019) or HNRNPC (Wu et al. 2018) can negatively regulate RLR signaling.

Surprisingly, we were not able to find expected innate immune dsRNA sensors of the RLR family, such as RIG-I or MDA5 using our approach. Our hypothesis is that their tight association to the substrate would impair the J2 antibody recognition and isolation of the long dsRNA. We did however isolate all four viral nsPs from SINV-infected cells suggesting that the DRIMS approach is suitable for the identification of proteins bound to the replication complex itself. Among the proteins isolated by DRIMS in SINV conditions, we also enriched for cytoplasmic, stress granules associated proteins such as G3BP1 and G3BP2, which have been shown to interact with either SINV nsP3 (Frolova et al. 2006) or nsP4 (Cristea et al. 2010) and play a role in the formation and function of SINV replication complexes. Moreover, G3BP1 was also shown to directly bind to viral dsRNA and poly(I:C) *in vitro* (Kim et al. 2019).

With our approach, we also isolated several RBPs with a known localization within nuclear paraspeckles. Paraspeckles are dynamic ribonucleoprotein (RNP) nuclear bodies built by protein liquid-liquid phase transition and defined by the presence of the structural long non-coding (lnc)RNA NEAT1 (nuclear-enriched abundant transcript 1) and the NEAT1-binding paraspeckle core proteins SFPQ (Splicing factor, proline-glutamine rich), NONO (Non-POU domain-containing octamer-binding protein) and PSPC1 (Paraspeckle component 1). Of note, NONO and SFPQ are able to interact as a heterodimer within the paraspeckles (Knott et al. 2016). These granules can act by sequestering several other proteins (*e*.*g*. MATR3, RBM14…) and hyper-edited nuclear RNAs containing Alu inverted repeats (IRAlus) and by modulating their molecular function (Nakagawa et al. 2018; Zhang and Carmichael 2001). In this work, we focused on the ubiquitous multifunctional protein SFPQ, which is involved in many aspects of nucleic acid biology in the nucleus (*i*.*e*. genome stability and transcriptional regulation, pre-mRNA splicing, localization and 3’end RNA processing) (Yarosh et al. 2015) and in the cytoplasm (*i*.*e*. mRNA post-transcriptional regulation, RNA granule formation especially in neurons) (Lim et al. 2020). In contrast to other paraspeckle-related proteins, such as NONO which is strictly nuclear, SFPQ showed a partial localization in the cytoplasm of HCT116 cells. We observed that this subcellular localisation did not change upon infection. Nonetheless, SFPQ cytoplasmic foci specifically accumulated upon SINV infection, which could be compatible with a differential binding to specific partners, independently of NONO, and a direct association with viral RNA. While NONO depletion did not seem to affect SINV infection, we showed that SFPQ depletion had a significant impact on the virus, by reducing SINV gRNA levels, protein levels as well as viral production. This phenotype, reproducible in two independent cell types (HCT116 and SK-N-BE(2) cells), suggests a proviral role for SFPQ on SINV infection.

SFPQ has also been reported to play a role in the regulation of several other RNA viruses (Yarosh et al. 2015). SFPQ interacts with HIV viral RNA to favour its nuclear-cytoplasmic export (Kula et al. 2013) and it also increases the efficiency of influenza virus mRNA polyadenylation (Landeras-Bueno et al. 2011). Moreover, SFPQ proviral effect has been reported during Hepatitis delta virus (HDV) (Greco-Stewart et al. 2006), human rhinovirus A16 (HRV16) (Flather et al. 2018), encephalomyocarditis virus (EMCV) (Zhou et al. 2019) and Severe acute respiratory syndrome coronavirus 2 (SARS-CoV-2) (Labeau et al. 2022) infections. Interestingly, SFPQ was also retrieved in previous proteomic approaches very different from ours to isolate proteins associated to either SINV or Semliki Forest Virus (SFV) (LaPointe et al. 2018; Varjak et al. 2013). Although SFPQ effect on viral infection was not characterized in these studies, these data support the implication of this protein on SINV and potentially other members of the same family.

Here, we experimentally validated the interaction of SFPQ with dsRNA and with the viral genomic and antigenomic RNAs. To our knowledge, this direct link between paraspeckle proteins and their binding to dsRNA has not been investigated so far. We also demonstrated that overexpression of a wild-type but not of an RNA binding SFPQ mutant increased viral capsid levels and viral titers in HCT116 cells treated with either siCTRL or siSFPQ against the 3’UTR (and therefore targeting specifically the endogenous transcript). These results indicate that the SFPQ RNA binding domains (RRM1-2) are necessary for the protein proviral function. However, our data indicate that overexpression of a myc-SFPQ WT in cells depleted of the endogenous protein (siSFPQ 3’UTR + myc-SFPQ WT) did not completely rescue the viral production to that of control cells (siCTRL + myc-BFP). This involves an additional level of regulation linking SFPQ to the regulation of SINV infection.

Interestingly, it is known that SFPQ can act as a transcriptional repressor of immune related genes. Virus-induced increase in paraspeckles size or number is linked with the re-localization of SFPQ to paraspeckles, leading to the transcriptional activation of the antiviral gene program (Imamura et al. 2014). We can therefore hypothesize that SFPQ depletion could affect viral replication and production in two ways: on one hand by decreasing the direct binding of cytoplasmic SFPQ to viral RNA, and on the other hand by causing the induction of antiviral immune genes. It is also possible that SFPQ binds to cellular RNAs that could be involved in the antiviral response, which could account for its proviral activity during SINV infection.

All in all, our results indicate that proteins once thought to be nuclear could have some unexpected functions in the context of viral infection. Accordingly, the characterization of the whole proteome associating with viral replication intermediates for different viruses and cell types might open new avenues of research in the field of antiviral immunity.

## Materials and Methods

### Cell culture, viral stocks and virus infection

Cell lines were maintained at 37°C in a humidified atmosphere containing 5% CO_2_. HCT116, BHK-21 and Vero cells were cultured in Dulbecco’s Modified Eagle medium (DMEM; Invitrogen) supplemented with 10% FBS. SK-N-BE(2) cells (95011815, Sigma-Aldrich) were maintained in 1:1 medium composed of Ham’s F12 medium (Gibco, Thermo Fisher Scientific Inc.) supplemented with 15% FBS and Eagle’s Minimal Essential Medium (ENEM) supplemented with 1% NEAA (Gibco, Thermo Fisher Scientific Inc.). SK-N-BE(2) cell differentiation was induced by 10 μM retinoic acid (RA) treatment (R2625, Sigma-Aldrich) for 5 days. Plasmids carrying a green fluorescent protein (GFP)-SINV genomic sequence or wild-type SINV genomic sequence (kindly provided by Dr Carla Saleh, Institut Pasteur, Paris, France) were linearized and used as a substrate for *in vitro* transcription using mMESSAGE mMACHINE capped RNA transcription kit (Ambion, Thermo Fisher Scientific Inc.) as in (López et al. 2020). Viral stocks were prepared in BHK-21 baby hamster kidney cells, and titers were measured by plaque assay in Vero cells. Cells were infected at a MOI of 10^−2^ or 10^−1^ as indicated in the figure legends and samples were harvested at 24 hours post-infection (hpi) unless specified otherwise.

### Standard plaque assay

Vero cells seeded either in 96- or 24-well plates were infected with viral supernatants prepared in cascade 10-fold dilutions for 1 h. Afterwards, the inoculum was removed and cells were cultured in 2.5% carboxymethyl cellulose for 72 h at 37ºC in a humidified atmosphere of 5% CO2. Plaques were counted manually under the microscope and viral titer was calculated according to the formula: PFU/mL = #plaques/ (Dilution*Volume of inoculum). PFU = plaque forming unit. For plaque visualization, the medium was removed, cells were fixed with 4% formaldehyde for 20 min and stained with 1x crystal violet solution (2% crystal violet (Sigma-Aldrich), 20% ethanol, 4% formaldehyde).

### DsRNA Immunoprecipitation and Mass Spectrometry (DRIMS)

Protein G Dynabeads were washed and resuspended in FA lysis buffer (1mM EDTA [pH 8.0], 50 mM HEPES-KOH [pH 7.5], 140 mM NaCl, 0.1% sodium deoxycholate (w/v), 1% triton X-100 (v/v), 1X protease inhibitors). Two micrograms (μg) of mouse anti-dsRNA J2 (SCICONS) or mouse anti-IgG (Cell signalling) were bound to 50 μL of beads for 1 h at room temperature. 80–90% confluent HCT116 cells in 10 cm^2^ plates were washed with cold PBS. Cells were scraped, transferred to a falcon and spun at 1000 rpm at 4°C for 5 min. For each IP, cell pellet from one 10 cm^2^ plate was lysed in 100 μL of Nuclei lysis buffer (50 mM Tris–HCl [pH 8.1], 10 mM EDTA [pH 8.0], 1% sodium dodecyl sulphate (SDS) (w/v), 1X protease inhibitor), incubated on ice for 10 min and diluted ten-fold with FA lysis buffer containing RNase inhibitor. Samples were sonicated in a BioRuptor (Diagenode) on high (“H”) for 5 min with 30 s on/off cycles. Following a spin at 13,000 rpm at 4°C for 5 min, supernatant was carefully transferred to a new tube and 40 μL of the volume were kept as input lysate. After 1 hour of pre-clearing at 4°C with yeast tRNA-blocked Protein G Dynabeads, Ab conjugated beads (50 μL) were added to the lysate and incubated for 2-3 hours at 4°C. Following magnetic separation, beads were washed twice with 1 mL of FA lysis buffer and twice with 1 mL of TE buffer (EDTA (10 mM, [pH 8.0]), Tris–HCl (100 mM, [pH 8.0]). Beads were incubated 10 min at 95°C with 40 μL 2X SDS Laemmli loading buffer (120 mM Tris/HCl [pH 6.8]; 20% glycerol; 4% SDS, 0.04% bromophenol blue; 10% β-mercaptoethanol) and proteins were analysed either by western blot or by silver staining after electrophoretic separation on polyacrylamide SDS-PAGE gel (Bio-Rad 4–20% Criterion™ TGX Stain-Free™ Protein Gel, 18 well, #5678094) using the SilverQuest™ Silver Staining Kit (ThermoFisher Scientific #LC6070) according to the manufacturer’s instructions.

For mass spectrometry experiments, proteins eluted from the beads were precipitated overnight with methanol containing 0.1% ammonium acetate and digested with trypsin after reduction and alkylation steps. Generated peptides were analysed by nanoLC-MS/MS on a QExactive□+□mass spectrometer coupled to an EASY-nanoLC-1000 (Thermo-Fisher Scientific, USA). The data were searched against a database containing the Human Swissprot sequences (2017_01), the SINV and GFP sequences with a decoy strategy. Peptides were identified with Mascot algorithm (version 2.5, Matrix Science, London, UK), and the data were imported into Proline 1.4 software [http://proline.profiproteomics.fr/]. The protein identifications were validated using the following settings: Mascot pretty rank□<□=□1, FDR□<□=□1% for PSM scores, FDR□<□=□1% for protein set scores. The total number of MS/MS fragmentation spectra was used to quantify each protein from three independent replicates.

Mass spectrometry data obtained for each sample, including the proteins identified by the Proline software suite and their associated spectral counts (SpC), were stored in a local MongoDB database and subsequently analysed through a Shiny Application built upon the R packages msmsEDA (Gregori J, Sanchez A, Villanueva J (2014). msmsEDA: Exploratory Data Analysis of LC-MS/MS data by spectral counts. R/Bioconductor package version 1.22.0) and msmsTests (Gregori J, Sanchez A, Villanueva J (2013). msmsTests: LC-MS/MS Differential Expression Tests. R/Bioconductor package version 1.22.0). Exploratory data analyses of LC-MS/MS data were thus conducted and differential expression tests were performed using a negative binomial regression model. In the edgeR functions used by the msmsTests package, an average prior count is added to each observation to shrink the estimated log-fold changes towards zero. Hence, infinite log-fold changes, that would arise from proteins with a null spectral count in one of the compared conditions, are avoided. The p-values were adjusted with FDR control by the Benjamini-Hochberg method and the following criteria were used to define differentially expressed proteins: an adjusted p-value < 0.05, a minimum of 5 SpC in the most abundant condition, and a minimum absolute Log2 fold change of 1. GO term analysis was performed using the EnrichR web-based tool (http://amp.pharm.mssm.edu/Enrichr) (Chen et al. 2013; Kuleshov et al. 2016). The direct interaction network for the enriched proteins was generated with the Cytoscape StringApp (Doncheva et al. 2019).

### SFPQ RNA Immunoprecipitation

Four million of either mock or SINV-infected HCT116 cells (MOI 0.1) were lysed 24 hpi using RIP immunoprecipitation buffer (50 mM Tris-HCl [pH 7.5], 150 mM NaCl, 5 mM EDTA, 0.05% SDS, 1% triton, 1 tablet of commercial protease inhibitor cocktail (Sigma Aldrich)). Lysates were treated for 20 min at 37°C with 1 μL of RNase-free DNase I by adjusting final concentration of MgCl_2_ (10 mM) and CaCl_2_ (5 mM). 1 μL of RNase inhibitor (Ribolock, ThermoFisher Scientific) was also added. Lysates were cleared by maximal centrifugation 10 min at 4°C. Supernatants were transferred in new Eppendorf tubes. Lysates were then precleared 1 h at room temperature with magnetic beads treated with tRNA solution (RIP buffer + 10 nM of tRNA). Lysates were incubated overnight with RIP immunoprecipitation buffer containing magnetic Protein G beads conjugated with human SFPQ/PSF antibody (sc-271796, Santa Cruz, Cliniscience) or negative control mouse IgG (5415S, Cell signalling, Ozyme). Beads were washed 3 times with RIP immunoprecipitation buffer, 3 times with wash buffer (50 mM Tris-HCl [pH 7.5], 200 mM NaCl, 5 mM EDTA, 0.05% SDS, 1% triton, 1 tablet of commercial protease inhibitor cocktail (Sigma Aldrich)) and twice with cold PBS 1X. Immunoprecipitated RNAs and proteins were isolated by TRIzol Reagent (Invitrogen) or Laemmli loading buffer, respectively. Extracted RNAs were analysed by RT-qPCR and proteins were analysed by western blot.

### Subcellular fractionation

Mock or SINV-infected (24 hours at MOI 0.1) HCT116 cells were washed with cold PBS 1X and gently resuspended in 3mL of hypotonic buffer (10 mM HEPES [pH 7.9], 1.5 mM MgCl_2_, 10 mM KCl, 1 tablet of commercial protease inhibitor cocktail (Sigma Aldrich)) and incubated on ice for 2 min. NP40 (0.4% final concentration) was added on the cell suspension. Cytoplasmic and nuclear fractions were separated by a centrifugation step at 500 g. The cytoplasmic fraction was removed and kept at 4°C. The nuclear pellet was washed by 3 times centrifugation steps (2 min at 500g) with hypotonic buffer. The nuclear pellet was resuspended in 1 mL of lysis buffer (20 mM Tris-HCl [pH 8.0], 100 mM KCl, 10 mM KCl, 0.2 mM EDTA, 0.5 mM DTT, 5% glycerol and 1 tablet of commercial protease inhibitor cocktail (Sigma Aldrich)).

### Western blotting

For protein analysis on total lysates and IP, proteins were extracted by collecting cell lysates in RIPA buffer (50 mM Tris-HCl [pH 7.5], 150 mM NaCl, 5 mM EDTA, 0.05% SDS, 1% triton, 1 tablet of commercial protease inhibitor cocktail (Sigma Aldrich)). Lysates were cleared by centrifugation at 13 000 rpm for 30 min at 4°C to remove cell debris and the supernatant was kept for western blotting. According to the experiment, 20 μg of proteins of each sample or both input and eluates from the IPs were heated in Laemmli loading buffer at 95°C for 5 min and loaded on a SDS-polyacrylamide electrophoresis gel (Bio-Rad 4–20% Mini-PROTEAN® TGX Stain-Free™ Protein Gels, 10 well, #4568093). Proteins were transferred to a nitrocellulose membrane by wet transfer in 1x Tris-Glycine + 20% ethanol buffer. Membranes were blocked in 5% milk diluted in PBS-Tween 0.2 % and probed with the antibodies indicated below (between 1:200 and 1:5000) in the same buffer. Viral proteins were detected using primary polyclonal antibodies against SINV capsid (kind gift from Dr Diane Griffin, Johns Hopkins University School of Medicine, Baltimore, MD). Anti-mouse-HRP (NXA931, GE Healthcare Thermo Fisher Scientific Inc.) or anti-rabbit-HRP (NA9340, GE Healthcare, Thermo Fisher Scientific Inc.) secondary antibodies were used (1:10000). Detection was performed using SuperSignal West Femto Maximum Sensitivity Substrate (Pierce, ThermoFisher Scientific) and visualized on a Fusion FX imaging system (Vilber).

### Immunostaining

HCT116 cells were plated on Millicell EZ 8-well slide (Merck Millipore) (60000 cells/well) and fixed with 4% formaldehyde (Sigma Aldrich), diluted in PBS 1X for 10 min at room temperature and incubated in blocking buffer (0.1% Triton X-100; PBS 1 X; 5% normal goat serum) for 1 h. Primary antibodies (see list below) were diluted in blocking buffer between 1:200 and 1:000 and incubated overnight at 4°C. Between each step, cells were washed with PBS 1X-Triton 0.1%. Cells were then incubated 1 h at room temperature with either goat anti-mouse Alexa 594 (A11032, Invitrogen) or goat anti-rabbit Alexa 488 (A11008, Invitrogen) secondary antibodies diluted at 1:1000 in PBS 1X-Triton X-100 0.1%. Cell nuclei were stained with DAPI (Life Technologies, Thermo Fisher Scientific) diluted at 1:5000 in PBS 1X during 5 min. Slides were mounted with coverslip with Fluoromount-G mounting media (Southern Biotech) and observed by confocal microscopy (LSM780, Zeiss).

### Antibodies

| NAME | species | ref | provider |
| --- | --- | --- | --- |
| ADAR1 | mouse | sc-271854 | Santa Cruz |
| DHX9 | rabbit | ab26271 | Abcam |
| DICER | rabbit | A301-936A | Bethyl |
| GAPDH-HRP | mouse | ab9482 | Abcam |
| Histone H3 | rabbit | ab1791 | Abcam |
| hNRNPA1 | mouse | sc-32301 | Santa Cruz |
| IgG | mouse | 5415S | Cell signalling |
| J2 | mouse | J2 | SCICONS |
| MATRIN3 | mouse | sc-81318 | Santa Cruz |
| myc | mouse | ab32 | Abcam |
| NONO | mouse | sc-166702 | Santa Cruz |
| PACT | rabbit | ab75749 | Abcam |
| PAXILLIN | rabbit | sc-5574 | Santa Cruz |
| PKR human | rabbit | ab32506 | Abcam |
| PSPC1 | mouse | sc-374181 | Santa Cruz |
| SFPQ/PSF | mouse | sc-271796 | Santa Cruz |
| SFPQ/PSF | rabbit | ab39148 | Abcam |
| TUBULIN | mouse | T6557 | Sigma-Aldrich |

### RNA extraction and RT-qPCR

Total and immunoprecipitated RNAs were extracted using Tri-Reagent (Invitrogen, Fisher Scientific) according to the manufacturer’s instructions. For total RNA, 1 μg of RNA was reverse transcribed using SuperScript IV Vilo Master Mix (Invitrogen, #11756050) according to the manufacturer’s instructions.

Real-time-PCR was performed with Maxima SYBR Green qPCR Master Mix (Applied Biosystem, #4309155) at an annealing temperature of 60°C on a CFX96 touch Real-Time PCR machine (Biorad) using the following primers:

SINV genome FW: 5’-CCACTACGCAAGCAGAGACG-3’;

SINV genome RV: 5’-AGTGCCCAGGGCCTGTGTCCG-3’;

GAPDH FW: 5’-CTTTGGTATCGTGGAAGGACT-3’;

GAPDH RV: 5’-CCAGTGAGCTTCCCGTTCAG-3’.

Strand-specific RT-PCR was performed as in (López et al. 2020). Briefly a (-) strand-specific reverse transcription was performed with a plus-sense primer annealing to the 5’ region of SINV genome (nucleotides 1-42) (5’-ATTGACGGCGTAGTACACACTATTGAATCAAACAGCCGACCA-3’).

RT reaction mix was set up and cDNA products were then amplified by PCR (25 cycles) with specific antigenome forward (5’-CATTCTACGAGCCGGTGCGC-3’) and reverse (5’-TAGACGTAGACCCCCAGAGTC-3’) primers using the GoTaq DNA polymerase (Promega) and analysed on a 1.5% agarose gel for analysis.

### SiRNA transfection

Transfection complexes were prepared using ON-TARGETplus Human siRNA (Horizon), siRNAs against ADAR (L-008630-00), BCLAF1 (L-020734-00), CSNK2A1 (L-003475-00), DDX3X (L-006874-02), DDX5 (L-003774-00), DHX15 (L-011250-01), DHX30 (L-017196-01), DHX9 (L-009950-00), DICER1 (L-003483-00), ELAV1 (L-016006-00), FASN (L-003954-00), FUS (L-009497-00), G3BP1 (L-012099-00), G3BP2 (L-015329-01), HNRNPA1 (L-008221-00), HNRNPL (L-011293-01), HNRNPQ/ SYNCRIP (L-016218-00), IGF2BP2 (L-017705-00) ILF3 (L-012442-00), MATR3 (L-017382-00-0005), NONO (L-007756-01-0005), PACT/ PRKRA (L-006426-00), PARP1 (L-006656-03), PNPT1 (L-019454-01), PSPC1 (L-020596-01-0005), RBM14 (L-020144-00-0005), RBMX (L-011691-01-0005), SFPQ (L-006455-00-0005), SRSF1 (L-018672-01), STAU1 (L-011894-00), STRBP (L-015439-02), TARBP2 (L-017430-00-0005), THRAP3 (L-019907-00), YBX1 (L-010213-00), YTHDF2 (L-021009-02), ZFR (L-019266-01), PKR (L-003527-00-0005), non-targeting Control Pool (D-001810-10-05) or siRNA against SINV 3’UTR (Integrated DNA Technologies) (López et al. 2020) at a final concentration of 50 nM and Lipofectamine 2000 transfection reagent (Thermo Fisher Scientific, #11668019) diluted in Opti-MEM transfection medium according to the manufacturer’s instructions and they were added to 20000 HCT116 cells in 96 well plates. Two days post-transfection, cells were subjected to SINV-GFP viral infection at MOI 10^−1^ for 24 hours before fixation and staining with Hoechst 3342 (labelling the nucleus). GFP quantification was carried out with the INCell Analyzer 1000 Cellular Imaging and Analysis System (GE Healthcare).

Alternatively, 600000 HCT116 cells were reverse transfected with 20 nM siRNA SFPQ or CTRL in 6-well plates, then transfected again 24 hours later and subsequently incubated for 24 hours before being infected with SINV-GFP at MOI 10^−1^ for 24 hours. For knock-down experiments in SK-N-BE(2) cells, 20000 cells were first transfected with 50 nM of siRNAs specific to SFPQ or CTRL in 6-well plates and 6 hours later, differentiation was induced by 10 μM retinoic acid (RA) treatment (R2625, Sigma-Aldrich). Samples were then analysed by epifluorescence microscopy, western blot, RT-qPCR or plaque assay.

### SFPQ CRISPR/Cas9 knock-out

Guide RNA (gRNA) sequences targeting human SFPQ were designed using the CRISPOR tool (http://crispor.tefor.net) and cloned into pX459-V2 vector (Addgene Plasmid #62988). Briefly, 100 ng of digested vector and 0.25 μM of the annealed oligos (guideRNA397FW: CACCGTCATCCTCCGTGATATCAGC; RV: AAACGCTGATATCACGGAGGATGAC; guideRNA64FW: CACCGTGTTAGTGGGTGCTGCCTAA; RV: AAACTTAGGCAGCACCCACTAACAC) were ligated and transformed into DH5 alpha cells. HCT116 cells were transfected with gRNA combined by pair. 24 hours after, the cells were treated with 1 μg/mL puromycin and the surviving cells were diluted to obtain 0.5 cells/ well in 96 well plates. Bulk cells were kept for validation. Two weeks later, cellular genomic DNA was extracted from individual colonies. Cells were lysed in 50 mM Tris-HCl [pH 8.0]; 100 mM EDTA [pH 8.0], 100 mM NaCl, 1% SDS and 0.1 mg of proteinase K, and incubated overnight at 55°C. Then, 50 ng of genomic DNA were amplified with GoTaq (Promega) using primers (IDT) to detect the deletion: SFPQ FW: TGGGTGTATCATCCAGTTCGG; SFPQ RV: GCAGAGGTCTCTGGTGTTTGAA. PCR reactions were loaded on a 1.5% agarose gel for analysis. Wild type genomic DNA and bulk were used as controls. SFPQ amplicons were gel purified and sequenced.

### MTT assay

MTT (3-(4, 5-dimethyl-2-thiazolyl)-2, 5-diphenyl-2-H-tetrazolium bromide) assay (Invitrogen) was performed to follow the proliferation of KO-SFPQ or siSFPQ HCT116 cells compared to WT or siCtrl HCT116 cells respectively. The cells were grown in 96-well plates and cultured for 24, 48 and 72 hours. In the knock-down assay, the cells were reverse transfected with 20nM of siRNA and transfected again after 24 hours with 20 nM of siRNA. 20 μL MTT (5 mg/mL, prepared in PBS1X) was added to each well containing 180 μL of DMEM and incubation was done during 3 h at 37°C, 5% CO2. Then the supernatant was removed and 100 μL DMSO were added into each well to dissolve the formazan crystals. The absorbance at 595 nm was measured with a microplate reader (SAFAS) after 30 min.

### SFPQ overexpression

The pCMV-myc-SFPQ wild-type (WT) (Addgene Plasmid #35183)(Rosonina et al. 2005) was used as a template to generate the pCMV-myc-SFPQ ΔRRM1-2 by mutagenesis using the InFusion HD kit from Clontech (catalog number 639649, Mountain View, CA), with the following primers:

ΔRRM1-2 FW: CTTACACACAGCGAACTCCTCGTCCAGTCATTGTGG; ΔRRM1-2 RV: GACTGGACGAGGAGTTCGCTGTGTGTAAGTTTTTCTC.

pCMV-myc-BFP was generated by deleting the SFPQ sequence from the CMV-myc-SFPQ wild-type (WT) and adding HindIII site by PCR using the following primers: HindIII pCMV-myc-PSF Rv:AAGCTTcccCAGATCCTCTTCTGAGATG; HindIII pCMV-myc-PSF Fw: gggAAGCTTCTGATATCGAATTCCTGCAG.

Then, the BFP insert was amplified from the pKLV-U6gRNA (BbsI)-PGKpuro2ABFP (Addgene Plasmid #50946) (Koike-Yusa et al. 2014) with the following primers: HindIII-TagBFP fw: gggAAGCTTacccgcaagcccggtgccgg; HindIII-TagBFP rv: cccAAGCTTtcaattGagcttgtgccccagtttgc, and it was ligated to the linearized vector.

For the plasmid overexpression experiment upon *SFPQ* KD with siSFPQ 3’UTR or siCTRL, the following conditions have been used: briefly, 2 × 10^5^ HCT116 cells were reverse-tranfected with 20nM of siRNA siCTRL or siSFPQ-3’UTR(Horizon discovery) in 12-well plates using the Lipofectamine 2000 transfection reagent (Thermo Fisher Scientific, #11668019) diluted in Opti-MEM transfection medium according to the manufacturer’s instructions and incubated for 24 hours at 37 °C, 5% CO2. Then, the cells were transfected with 1μg of of either the pCMV-myc-SFPQ WT or ΔRRM1-2, or a control (pCMV-BFP) using the Lipofectamine 2000 and incubated again for 24 hours before being infected with SINV-GFP at MOI 0.1 for 24 hours. Proteins lysates and supernatants were harvested and the samples were analysed by western and plaque assay, respectively. Mock-infected cells were also collected for protein analysis.

## Data availability

The mass spectrometry proteomics data have been deposited to the ProteomeXchange Consortium via the PRIDE partner repository (Perez-Riverol et al. 2019) with the dataset identifier PXD024554

## Acknowledgments

The authors would like to thank all members of the Pfeffer laboratory for discussion, Amélie Weiss for technical help with GFP quantification with the INCell Analyzer 1000 Cellular Imaging System, Dr. Olivier Petitjean for cloning the pCMV-mycBFP construct, Dr Carla Saleh for providing the SINV WT and GFP clones and Dr Diane Griffin, for providing CAPSID and nsP2 antibodies. This work was funded by the European Research Council (ERC-CoG-647455 RegulRNA) and was performed under the framework of the LABEX: ANR-10-LABX-0036_NETRNA and ANR-17-EURE-0023, which benefits from a funding from the state managed by the French National Research Agency as part of the Investments for the future program. This work has also received funding from the People Programme (Marie Curie Actions) of the European Union’s Seventh Framework Program (FP7/2007-2013) under REA grant agreement n° PCOFUND-GA-2013-609102, through the PRESTIGE program coordinated by Campus France (to EG). The mass spectrometry instrumentation was funded by the University of Strasbourg, IdEx “Equipement mi-lourd” 2015.

## Author contributions

SP and EG conceived the project. SP and EG designed the work and analysed the results. EG, MM, PL, AF, JC performed the experiments. EG set up the DRIMS experiments and performed the J2-IP western blots and dot blots. JC performed the bioinformatic analysis of the proteomic data, BCWM developed the MongoDB database and the Shiny application used to analyse the mass spectrometry data, EG performed the GO-enrichment and STRING analysis. EG performed the siRNA screen. EG and MM performed the SFPQ knock-down experiments, the western blot and qPCR analysis, EG and PL performed the plaque assays. EG and MM performed the immunofluorescence and data acquisition, MM performed the image analysis. MM generated the SFPQ +/-cell line and set up the SFPQ-RIP, PL and EG performed and analysed SFPQ KD and SINV-GFP infections in differentiated SK-N-BE(2) cells. MM generated the SFPQ mutant plasmid, MM and EG performed the rescue experiments. PH coordinated the proteomic experiments. EG and MM drafted the manuscript and designed the figures. EG and SP wrote the manuscript with input from the other authors. SP and EG coordinated the work. SP assured funding. All authors revised the final manuscript.

## Supplementary figures and tables legends

**Table S1**. Details of the differential expression analysis results obtained for the enriched proteins in SINV-J2-IP *versus* mock-J2-IP. Proteins are listed based on increasing adj. pvalue.

**Table S2**. Details of the differential expression analysis results obtained for the enriched proteins in SINV-J2-IP *versus* SINV-IgG-IP. Proteins are listed based on increasing adj. pvalue.

**Table S3**. Details of the differential expression analysis results obtained for the enriched proteins in mock-J2-IP *versus* mock-IgG-IP. Proteins are listed based on increasing adj. pvalue.

**Figure S1. DsRNA-associated protein profiles in mock and SINV-GFP infected HCT116 cells A)** Silver-stained 4-20% Tris-SDS-PAGE gel of the total lysates (INPUT) and eluates from J2- or IgG control-IP in mock and SINV-infected cells used for proteomic analysis. **B)** Hierarchical clustering of the J2-IP SINV (in orange) and mock (in green) infected samples. **C-D)** Volcano plots of dsRNA-IP (J2-IP) *versus* IgG control-IP **(C)** in SINV-infected and **(D)** in mock-infected conditions using data from three replicates. Red and blue dots represent proteins which are significantly enriched or depleted (adjusted p-value < 0.05, a minimum of 5 SpC in the most abundant condition and abs(Log2FC) > 1), respectively, in the J2-IP samples compared to the IgG control-IP ones. Viral non-structural proteins are indicated in orange. Known dsRNA binding proteins are indicated by a black square. Proteins present in paraspeckles are indicated by a black circle and a light blue name. **E)** GO term enrichment analysis of the dsRNA-associated proteins overrepresented in SINV-J2-IP compared to mock-J2-IP using the Enrichr software. Enriched GO terms of molecular functions (green), cellular components (yellow) and biological processes (purple) are respectively sorted by p-value.

**Figure S2. SFPQ depletion by siRNA in HCT116 and SK-N-BE(2) cells reduces SINV-GFP infection. A)** MTT assay results on HCT116 cell proliferation upon treatment with siCTRL (in blue) or siSFPQ (in orange), measured at 24, 48 and 72 h of growth. **B-C**) Western blot performed on samples from siCTRL and siSFPQ-treated HCT116 cells in mock and SINV infection at the indicated time points and MOI. Antibodies directed against SFPQ and the viral capsid protein were used. Tubulin was used as loading control. **D**) Representative pictures of SINV-GFP infected cells in siCTRL and siSFPQ treated SK-N-BE(2) cells. GFP expression was measured by microscopy. BF, brightfield. Scale bar: 100 μm. **E**) Western blot performed on non-transfected (NT) mock and infected cells and on SINV-GFP infected cells upon siCTRL and siSFPQ treatment. Antibodies directed against SFPQ and the viral capsid protein were used. Tubulin was used as loading control. **F)** Viral production of SINV-GFP upon siCTRL and siSFPQ transfection measured by plaque assay (PFU/mL) on three independent biological replicates. Infection conditions: 24 hours at MOI of 0.1. Each replicate is indicated with a different coloured dot. Error bars represent the mean +/-standard deviation (SD) of three independent experiments, * p < 0.05, paired Student’s t test.

**Figure S3. Depletion of one *SFPQ* allele by CRISPR/Cas9 in HCT116 cells reduces SINV-GFP infection. A)** Schematic representation of the CRISPR/Cas9 knockout strategy used to generate the (+/-) *SFPQ* HCT116 cells. Two gRNAs spanning the exon 2/intron 2 junction were used to direct Cas9 cleavage and the deletion of about 300 bp including the intron 2 5’splice site (5’ss). **B**) Agarose gel showing the PCR fragments corresponding to the WT and KO alleles respectively. Sequencing of the PCR fragment corresponding to the KO allele is shown in (A). **C-D-E)** Effect of *SFPQ* heterozygous knock-out in HCT116 cells. **C)** Representative pictures of SINV-GFP infected cells in WT and (+/-) *SFPQ* HCT116 cells. GFP expression was measured by microscopy. BF, brightfield. Scale bar: 100 μm. **D)** Western blot performed on mock and infected cells in WT and (+/-) *SFPQ* HCT116 cells. Antibodies directed against SFPQ and the viral capsid protein were used. Tubulin was used as loading control. **E)** Viral production of SINV-GFP in WT and (+/-) *SFPQ* HCT116 cells measured by plaque assay (PFU/mL) on three independent biological replicates. Infection conditions: 24 hours at MOI of 0.1. Each replicate is indicated with a different coloured dot. Error bars represent the mean +/-standard deviation (SD) of three independent experiments, * p < 0.05, paired Student’s t test. **F**) MTT assay results on cell proliferation of WT (in blue) and (+/-) *SFPQ* (in orange) HCT116 cells measured at 24, 48 and 72 h of growth.

**Figure S4. SFPQ localization and binding to SINV antigenomic RNA. A**) Confocal immunofluorescence analysis on HCT116 cells in mock and SINV WT infection conditions. Antibodies against G3BP1, NONO and SFPQ were used (yellow signals). Antibody against dsRNA (J2) was used as a positive control of infection. DAPI was used to stain the nuclei (in blue in the merge). Magnification 20X(4X). **B**) Western blot against SFPQ upon cell fractionation on HCT116 cells in mock and SINV-GFP infection conditions. Antibodies against Histone 3 (H3) and Paxillin were used to control the nuclear (N) and cytoplasmic © fractions respectively. GFP was used as a control of the infection. **C**) Confocal co-immunofluorescence analysis on mock and SINV WT infected HCT116 cells. Anti-SFPQ rabbit antibody (in purple) or anti-dsRNA J2 mouse antibody (in yellow) were used. Merge is shown in Figure 4B. DAPI was used to stain the nuclei (in grey). Magnification 63X(3X). **D**) Strand-specific RT-PCR on SINV antigenome using total RNA (INPUT) and RNA isolated upon IgG or SFPQ RIP from SINV-infected samples. RT-, negative control.

**Figure S5. SFPQ binding to viral RNA is independent of ssRNA. A)** RNA analysis on agarose gel upon RNase T1 treatment on mock and SINV infected total lysates. **B)** Anti-SFPQ western blot performed on total lysate (INPUT), J2- or IgG-IP in mock and SINV infected HCT116 cells with or without RNase T1 treatment. Antibodies against PKR (positive control, dsRBP) and AGO2 (negative control, ssRBP) were used.

**Figure S6**. Western blot in mock-infected HCT116 cells first treated with siCTRL or siSFPQ3’UTR and then transfected with a control myc-BFP, a myc-SFPQ WT or a myc-SFPQ ΔRRM1-2 plasmid. Antibodies directed against the myc-tag and the SFPQ protein were used. Ponceau staining and tubulin antibody were used as loading controls.

## Notes

### Competing Interest Statement

The authors have declared no competing interest.

### Summary of Updates

Addition of new data mainly the use of a RNA-binding mutant version of SFPQ to show the importance of RNA binding for the proviral phenotype of the protein. Modifications in the text.

