## Supplementary figures and images for "Proteomics-based determination of double stranded RNA interactome reveals known and new factors involved in Sindbis virus infection"

### Figure S1

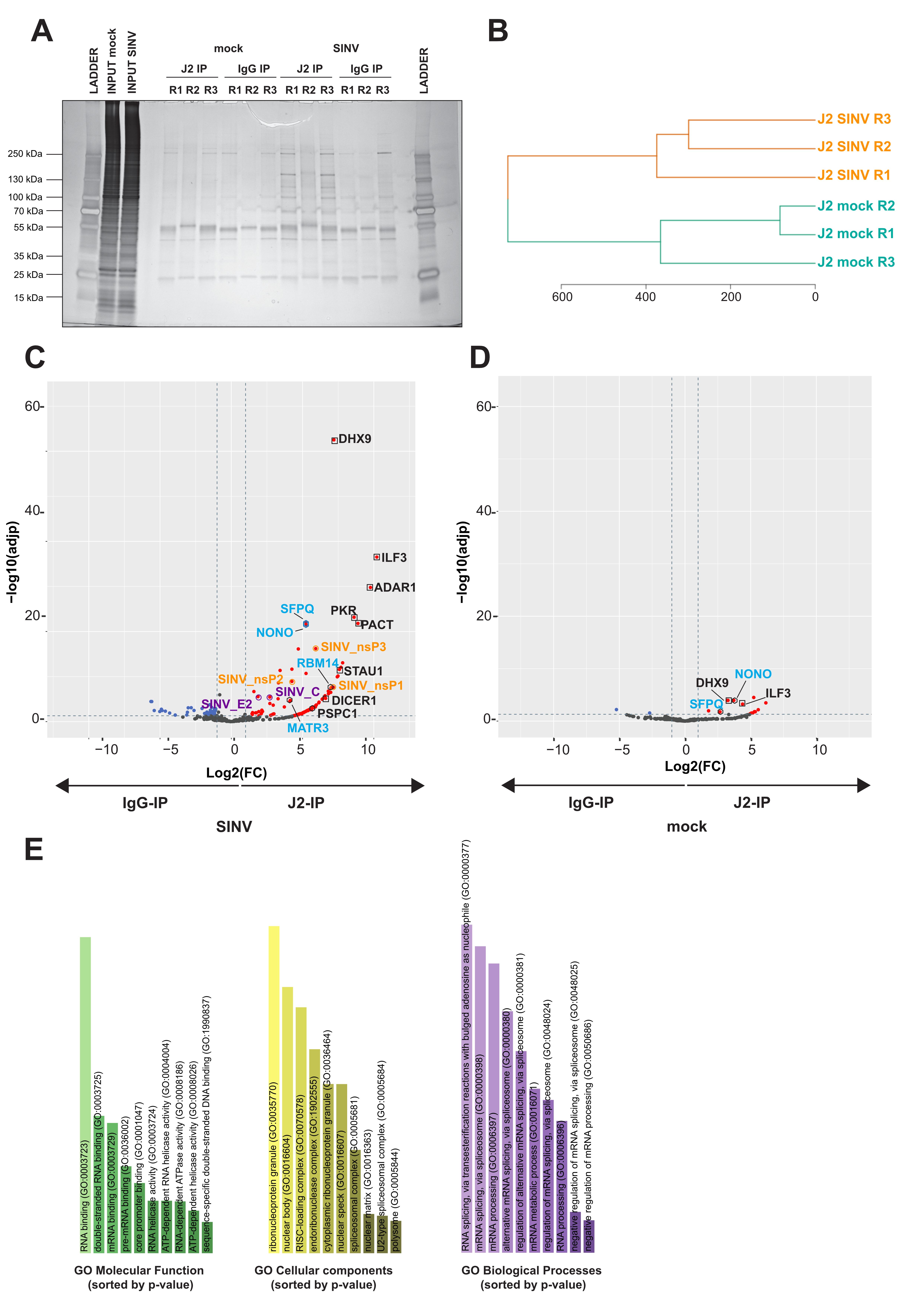

### Figure S2

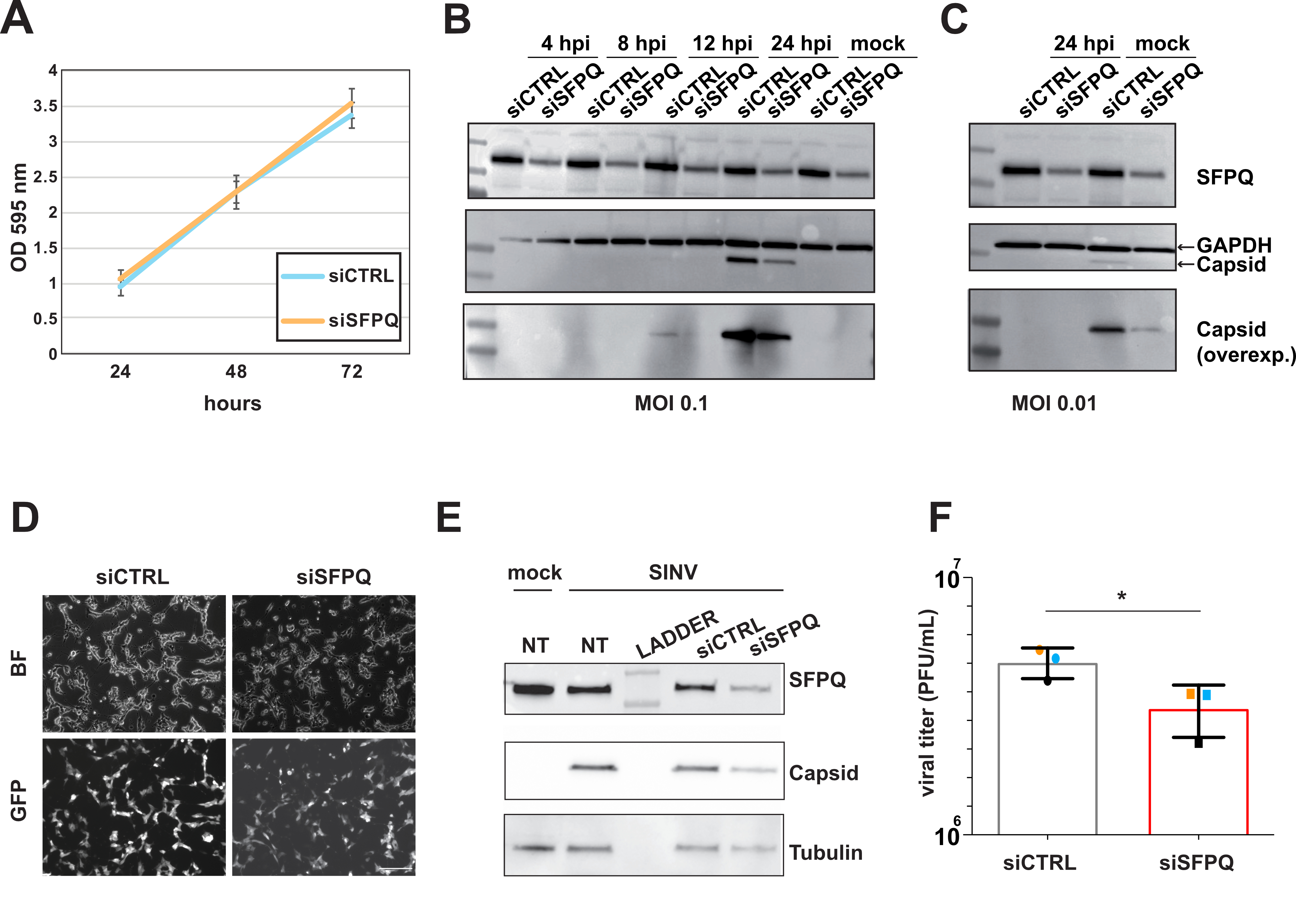

### Figure S3

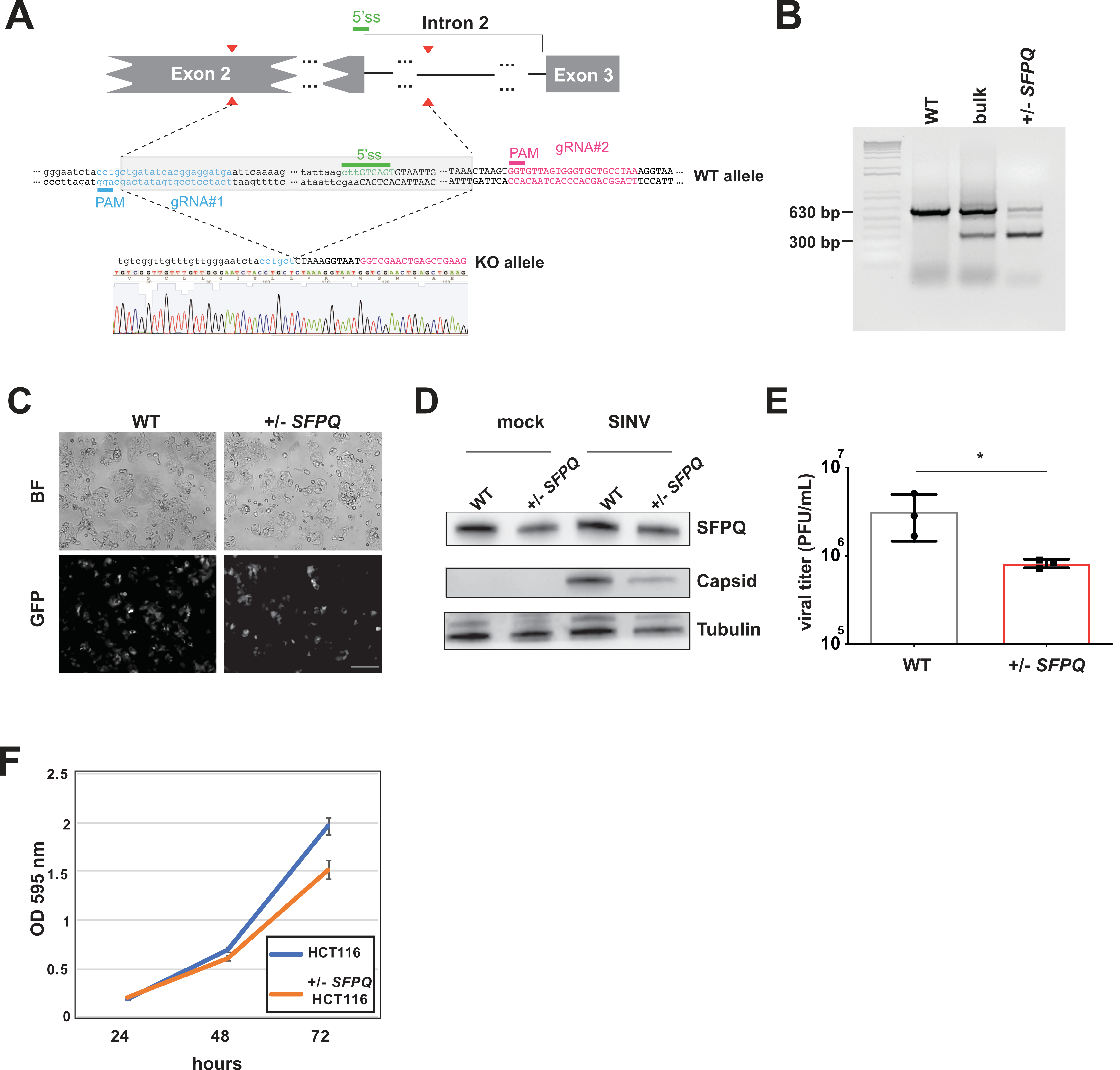

### Figure S4

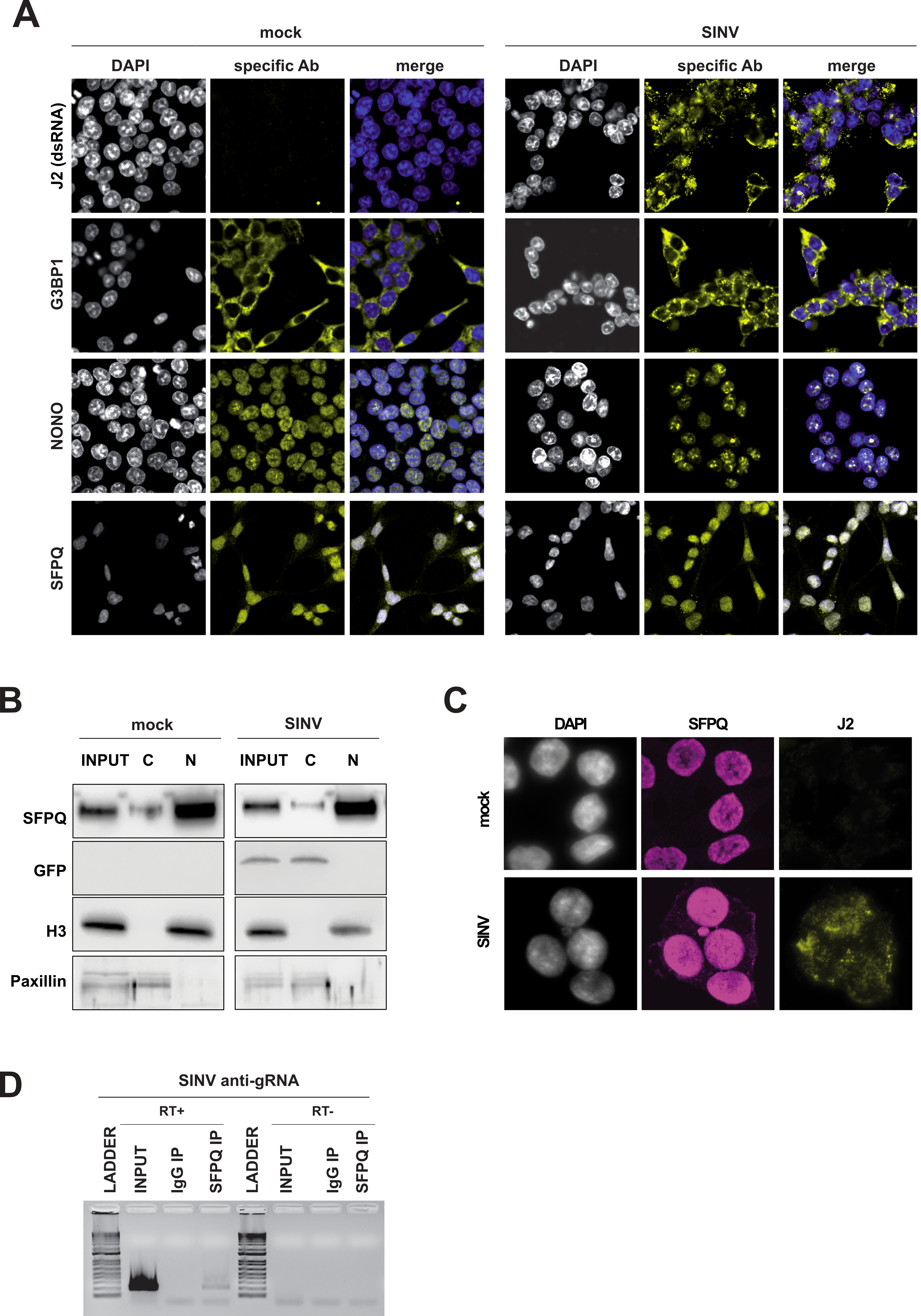

### Figure S5

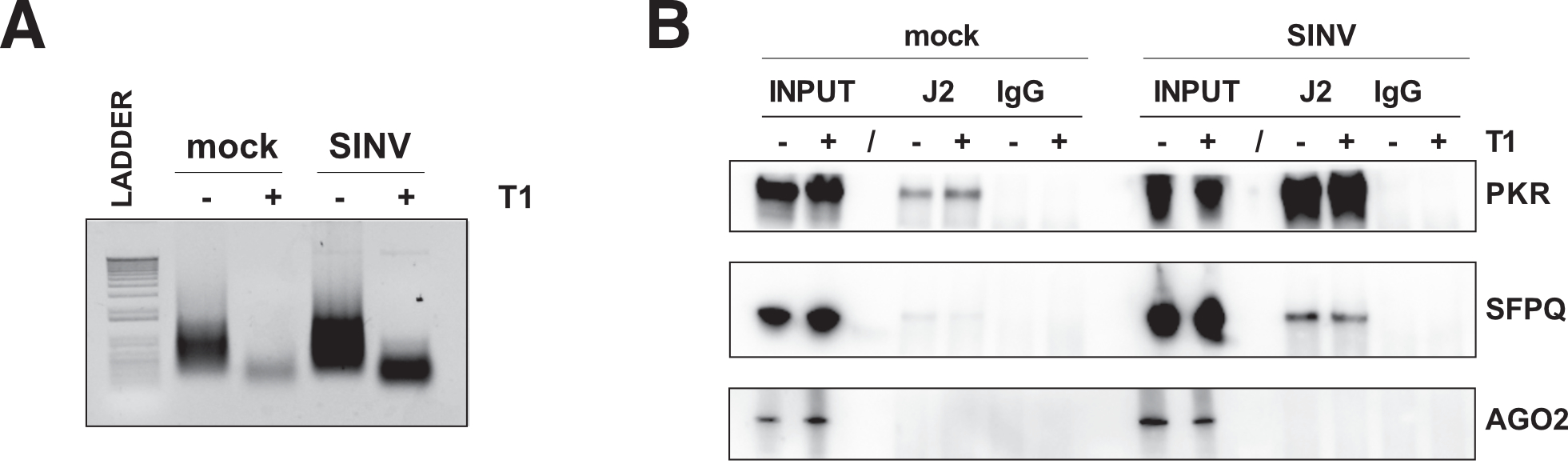

### Figure S6

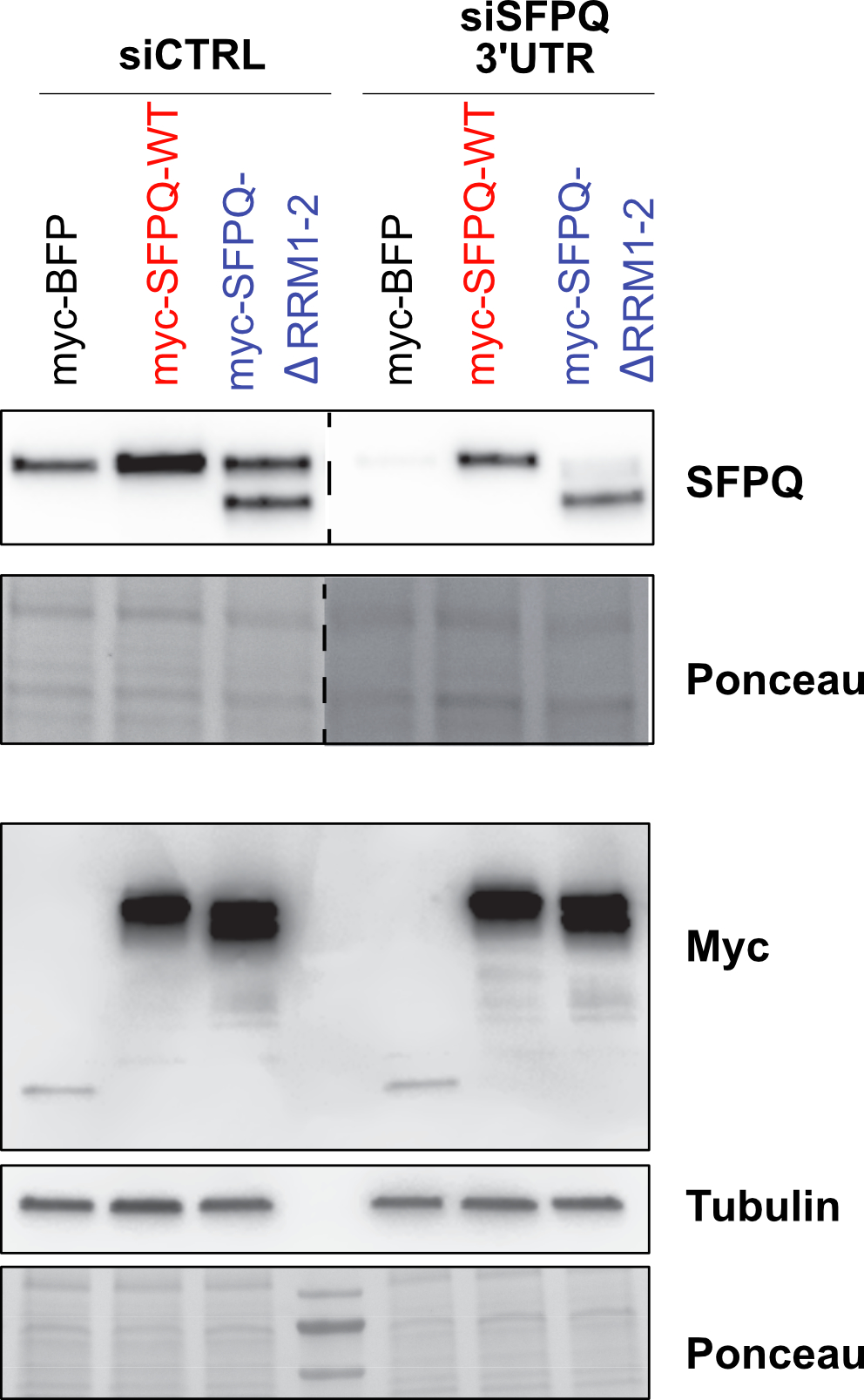
